# Small mammalian herbivores at moderate densities facilitate livestock growth by improving vegetation composition in grasslands

**DOI:** 10.64898/2026.04.06.716644

**Authors:** Zhiwei Zhong, Bingbo Ni, Douglas Lawton, Xiaofei Li, Xiaona Zheng, Huakun Zhou, Junhu Su, Wenjin Li, Fujiang Hou, Zhenggang Guo, Quanmin Dong, Shikui Dong, Christopher R. Dickman, Jens-Christian Svenning, Ying Gao, Zhibin Zhang

**Author notes:** These authors contributed equally to this work (Zhiwei Zhong, Bingbo Ni).

## Abstract

Small mammals and large herbivores have co-evolved in grasslands for millions of years, yet how they interplay remains unclear. Although large herbivores can significantly affect the smaller ones, the reversal effect is largely unknown. On the Qinghai–Tibetan Plateau, plateau pikas (*Ochotona curzoniae*) are often considered pests that compete with livestock at high densities. Using field experiments, we show that the presence of pikas facilitates weight gains of yaks (*Bos grunniens*) by improving vegetation composition at a moderate density level. Compared to the pika-present treatment, pika removal dramatically increased cover of the poisonous *Stellera* forbs by two-fold, reducing the abundance and protein content of palatable grasses and sedges, yak foraging efficiency, and yak weight gain by up to 42%. Notably, we found a humped relationship between yak weight gains and pika burrow densities in the pika-present plots; the facilitative effect of pikas on yaks was highest at about 200 burrows/ha, but shifted to a competitive effect at pika densities exceeding 400 burrows/ha. These results provide the first empirical evidence that maintaining a moderate density of small mammalian herbivores can benefit growth performance of livestock by improving vegetation composition. Our study highlights the significance of moderate populations of ecosystem-engineering small mammals in sustaining pastoral productivity in rangelands.

## Introduction

Both large and small mammalian herbivores are key consumers in grasslands worldwide, exerting profound impacts on ecosystem structure and functions (Dickman, 1999; Keesing, 2000; Davidson et al., 2012; Ripple et al., 2015; Pringle et al., 2023; Barbero-Palacios et al., 2024; Lundgren et al., 2024). They have co-evolved with grasslands and with each other over millions of years, but how these mammals co-exist remains poorly understood (Smith and Foggin, 1999; Detling, 2006; Davidson et al., 2010; Li et al., 2019; Speakman et al., 2021). By sharing food and other resources, guilds of different herbivore species can participate in varied interactions ranging from negative (competition) to positive (facilitation) for one or more of the species (Arsenault and Owen-Smith, 2002; Odadi et al., 2011; Augustine and Springer, 2013; Barrio et al., 2013; Zhong et al., 2014; Harris et al., 2015; Anderson et al., 2024). However, such interactions often are assumed to be highly asymmetric: large herbivores have been demonstrated to affect the abundance, diversity, and demography of small mammals (Keesing, 1998; Smit et al., 2001; Bakker et al., 2009; Munoz et al, 2009; Badingqiuying et al., 2018; Wang et al., 2019; Trepel et al., 2024), whereas the potential for reverse effects have received little attention (Detling 2006; Wang et al., 2020; Augustine and Derner, 2021; Augustine et al., 2024).

Small mammals, including pikas, voles, prairie dogs, pocket gophers, and other small herbivores, are not only primary consumers but also key ecosystem engineers in many grasslands (Jones et al., 1994; Smith and Foggin, 1999; Zhang et al., 2003; Davidson et al., 2012; Augustine et al., 2023). Through their feeding, clipping, and burrowing activities they profoundly alter vegetation and soil properties, with potential cascading effects on co-occurring large herbivores (Detling, 2006; Davidson et al., 2012; Wang et al., 2020; Augustine and Derner, 2021; Augustine et al., 2024). In rangelands and other less productive ecosystems, small herbivorous mammals are often considered to be pests because their population outbreaks can lead to competition with livestock for food (Zhang and Wang, 1998; Dickman, 1999; Singleton et al., 2010; Singleton et al., 1999). However, it remains unclear whether the competitive effects of outbreaks of small mammalian herbivores on livestock diminish or even shift to facilitation when the smaller species are at lower, non-outbreak levels as predicted by the ecological nonmonotonicity theory (Zhang et al., 2015). Livestock grazing currently uses ∼77% of global agricultural land, and sustains billions of people worldwide (Maestre et al., 2022). A critical assessment of small mammals’ impacts on livestock production is urgently needed to guide management for both production and biodiversity conservation.

The Qinghai-Tibetan Plateau supports approximately 14 million yaks (*Bos grunniens*), forming one of the world’ s most extensive pastoral systems (Long et al., 2008). These herds are vital to pastoral livelihoods and key ecosystem functions across the region’ s 2.5 million km² expanse (Dong et al., 2020). The dominant small mammal species—plateau pika (*Ochotona curzoniae*)—is an iconic keystone species that commonly coexists with yaks, and is the focus of long debate on whether its impact of yaks is positive or negative (Fan et al., 1998, 1999; Smith and Foggin, 1999; Delibes-Mateos et al., 2011). In regions with high population densities (e.g., over 500 active burrows/ha), plateau pikas can suppress livestock production by heavily consuming nearly all plant species foraged by yaks, leading to extensive poisoning campaigns targeted at eradicating these small mammals (Fan et al., 1998, 1999; Zhang et al., 1999). At low and moderate densities of pikas (e.g., below 200 active burrows/ha), however, competition for food is mitigated and dietary partitioning can occur: yaks graze selectively on monocotyledonous plants such as grasses and sedges (Pan et al., 2019; Li et al., 2023), whereas pikas prefer to clip and feed on the leaves of dicotyledonous plants (Jiang and Xia 1985; Liu et al., 2008). Thus, whether plateau pikas and yaks compete for pasture largely may depend on their diets and population densities.

In line with global trends in weed invasions in rangelands (DiTomaso, 2000), intensive livestock grazing and other human disturbances have facilitated grassland degradation on the Qinghai-Tibet Plateau, creating opportunities for the proliferation of toxic weeds (Lu et al., 2012). Among these, the *Stellera chamaejasmehas* become a predominant species, now covering approximately 20 million hectares, as it outcompetes palatable grasses and sedges while thriving under grazing pressure (Li and Zhao, 2025). The expansion of this poisonous plant species on the Qinghai-Tibetan Plateau poses a dual threat to livestock, either by causing direct toxicity (Keeler et al., 2013) or by reducing forage availability through competition with palatable species for light and soil nutrients (Lu et al., 2012; Li and Zhao, 2025). However, plateau pikas may indirectly benefit yak foraging by selectively clipping large plants (Liu et al., 2019; Zhang et al., 2020), especially those poisonous forbs such as *S. chamaejasme* (Fan et al., 1998) (Fig. 1)—a behavior that reduces predation risk (Dickman, 2022; Zhong et al., 2022) but also suppresses these competitors, potentially enhancing the quality and quantity of desirable grasses and sedges. Despite these plausible interactions, empirical studies directly testing such interactions remain scarce.

**Figure 1.**
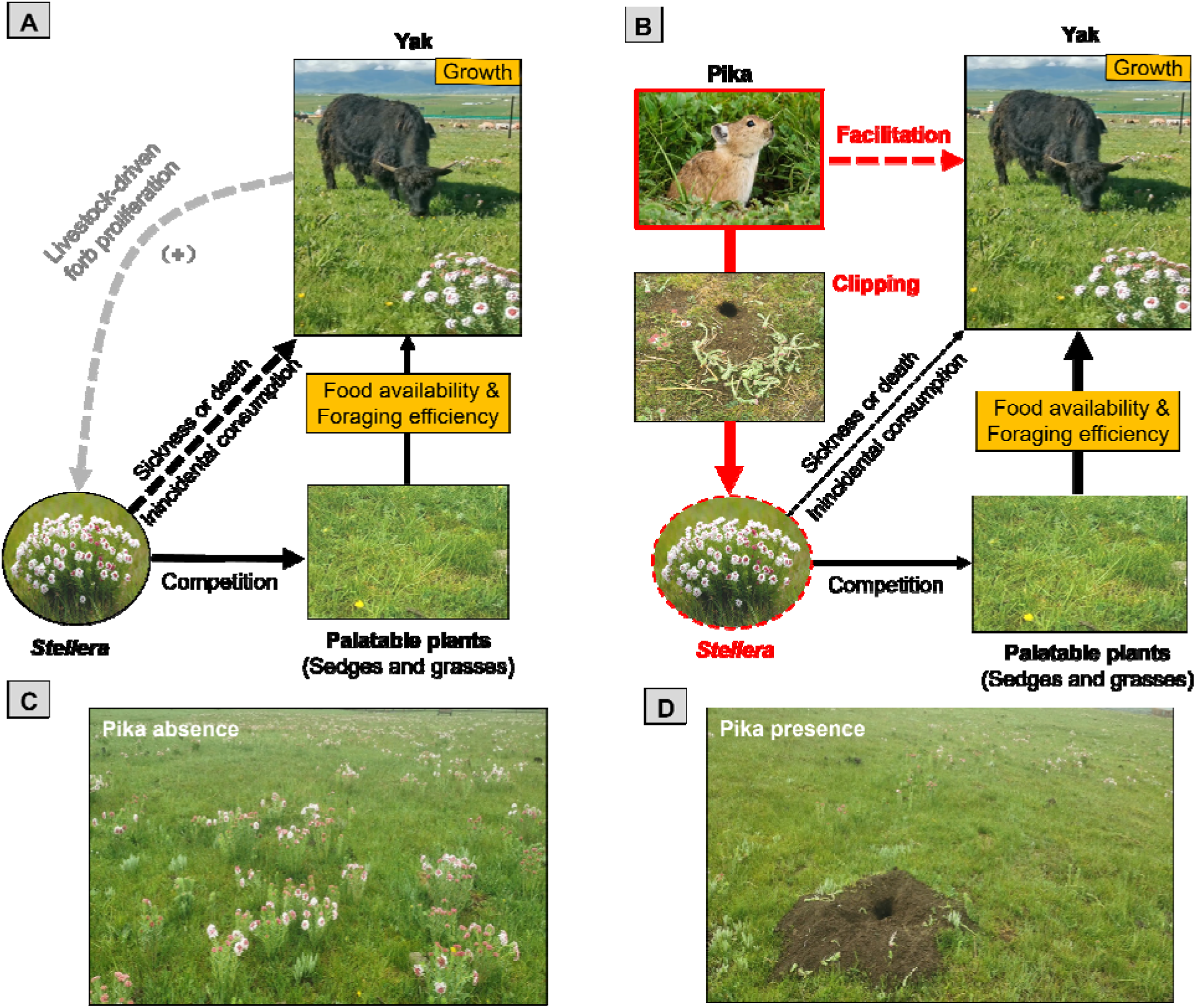
Conceptual diagram of the hypothesized facilitative effects of pikas (*Ochotona curzoniae*) on yak (*Bos grunniens*) growth performance on the Qinghai-Tibetan Plateau. (**A**) In the absence of pikas, the poisonous forb *Stellera chamaejasme* exerts strong negative effects on the growth of yaks by outcompeting palatable grasses and sedges, thereby reducing forage availability and foraging efficiency. (**B**) In the presence of pikas, clipping of *Stellera* suppresses its abundance, mitigating its negative impact and promoting yak growth. Grey dashed line and “+” indicate livestock grazing promotes the proliferation of the poisonous forbs by suppressing palatable grasses and sedges. Black dashed lines indicate negative effects of poisonous plants on yaks, black solid lines indicate competition between plant groups, and red solid lines indicate pika suppression of *Stellera*. Line thickness represents the relative strength of putative species interactions. (**C**, **D**) show pika clipping activity and its effect on *Stellera* abundance at the study site. Credits: Xiaona Zheng (photographs).

Here, we experimentally assess how a moderate population density of pikas (∼200 active burrows ha⁻ ¹) affects yak performance in an alpine ecosystem of the Qinghai–Tibetan Plateau. We hypothesized that, at moderate densities, plateau pikas facilitate yaks by suppressing tall poisonous forbs, thereby increasing the abundance and nutritional quality of palatable grasses and sedges during the growing season (June–August). Specifically, we predicted that (i) pika clipping reduces poisonous plant cover; (ii) this enhances the availability and quality of palatable forage (i.e., grasses and sedges) for yaks; (iii) yaks increase their foraging efficiency in the presence of pikas; and (iv) these cascading effects ultimately improve yak weight gain (Fig. 1). To test these predictions, we first conducted two field surveys to examine diet partitioning between pikas and yaks, and the associations among pika density, *Stellera* abundance, and yak grazing activity. We then performed an in-situ field experiment using 150 × 150 m fenced enclosures to test the interactive effects of pikas and *Stellera* on yak body growth and the underlying mechanisms driving these effects.

## Results

In the field surveys, we found that pikas and yaks have distinct diets: pikas very frequently clipped but did not eat large, poisonous *Stellera* plants and fed mostly on other forbs, whereas yaks consumed greater proportions of grasses and sedges (Fig. 2A,B, Table S1, S2), supporting findings of previous studies (Jiang and Xia, 1985; Fan et al., 1998; Liu et al., 2008; Pan et al., 2019; Li et al., 2023). We also found that the cover abundance of *Stellera* was associated negatively with the density of active pika burrows (R^2^ = 0.39, *P* <0.001; Fig. 2C, Table S3) and with yak foraging activity, as indicated by dung density (R^2^ = 0.43, *P* < 0.001; Fig. 2D, Table S4).

**Figure 2.**
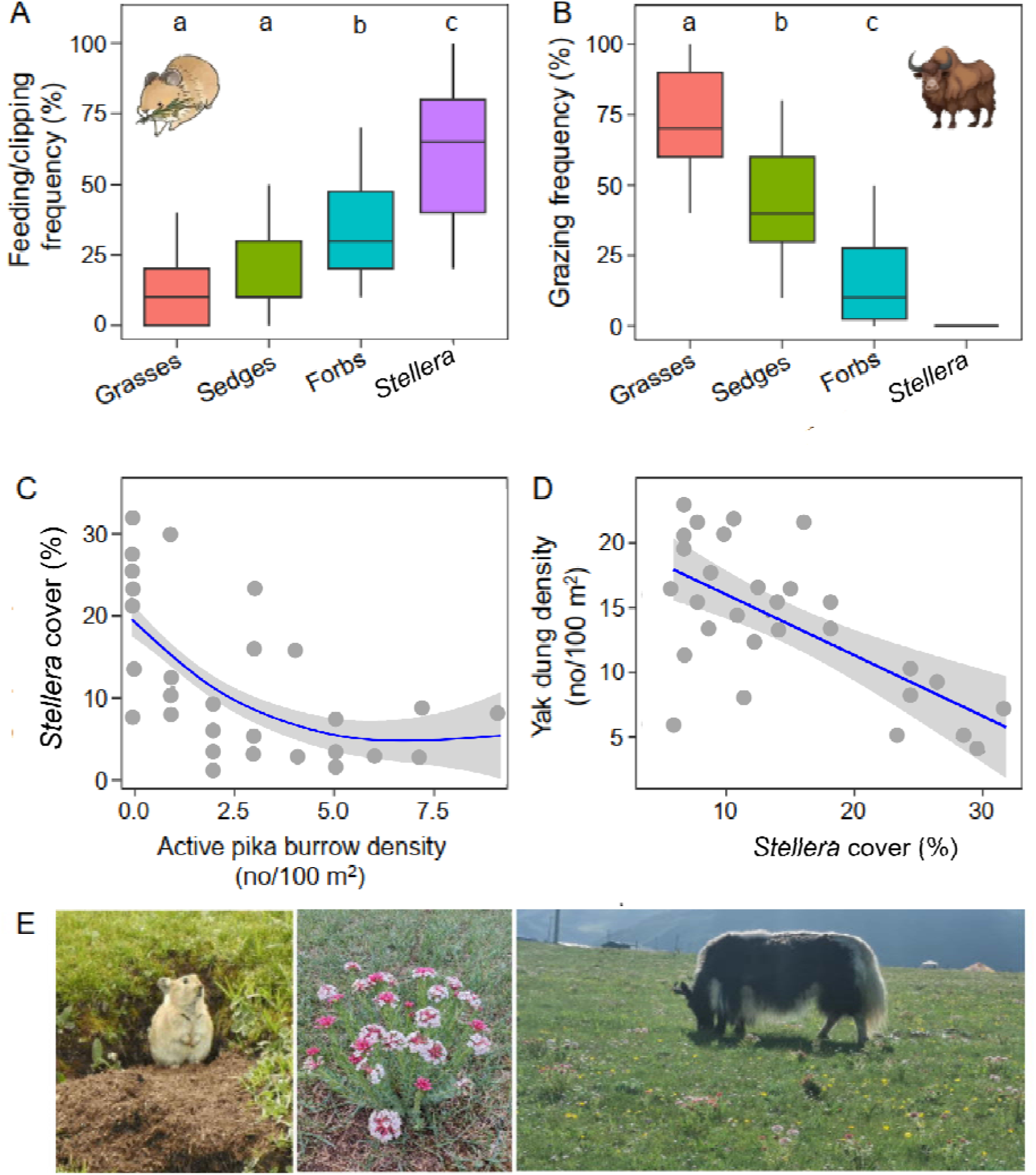
Diet selection of pikas and yaks and their potential interactions mediated by poisonous plants based on field surveys in July 2021. (**A**) Feeding and clipping frequencies of pikas, and (**B**) grazing frequencies of yaks on grasses, sedges, forbs, and *Stellera* across 10 2 × 2 m plots and 10 250 m transects, respectively. (**C**) Relationship between active pika burrow density and *Stellera* cover, and (**D**) between *Stellera* cover and an index of yak grazing activity (dung density) in 30 10 × 10 m plots. (**E**) Photographs showing (left to right) a pika at its burrow entrance, flowering *Stellera*, and a yak grazing among *Stellera* plants. Different letters above bars indicate significant differences at *P* < 0.05. Credits: Xiaona Zheng (photographs).

In the manipulative field experiment, pika removal led to a 90% reduction in the number of active burrows per hectare, with 182.9 ± 24.5 burrows/ha in pika-present plots compared to 19.0 ± 1.6 burrows/ha in the pika-absent plots (Fig. 3A, Table S5). In the presence of *Stellera*, pika absence reduced daily weight gains of yaks by 42% compared to those in the pika-present treatment, but such effects disappeared once *Stellera* was removed (Fig. 3B, Table S5, S6). Notably, yak weight gains exhibited a hump-shaped relationship with pika densities, with the highest growth at about 200 active burrows/ha, after which, yak growth decreased linearly as pika density increased (R^2^ = 0.75, *P* < 0.001; Fig. 3C, Table S7). Weight gain was ∼ 0.3 kg/yak/day at zero pika density, peaked at ∼0.4 kg/yak/day around 200 burrows/ha, and fell below the baseline when pika density exceeded 400 burrows/ha. These data indicate that pikas facilitated yak weight gains by up to 30% across 0-400 burrows/ha, with maximum benefit at ∼ 200 burrows/ha, whereas higher pika densities (> 400 burrows/ha) suppressed yak weight gains. The pika-yak facilitation peak and the facilitation-competition transition therefore occur at approximately 200 and 400 burrows/ha of pika density levels, respectively.

**Figure 3.**
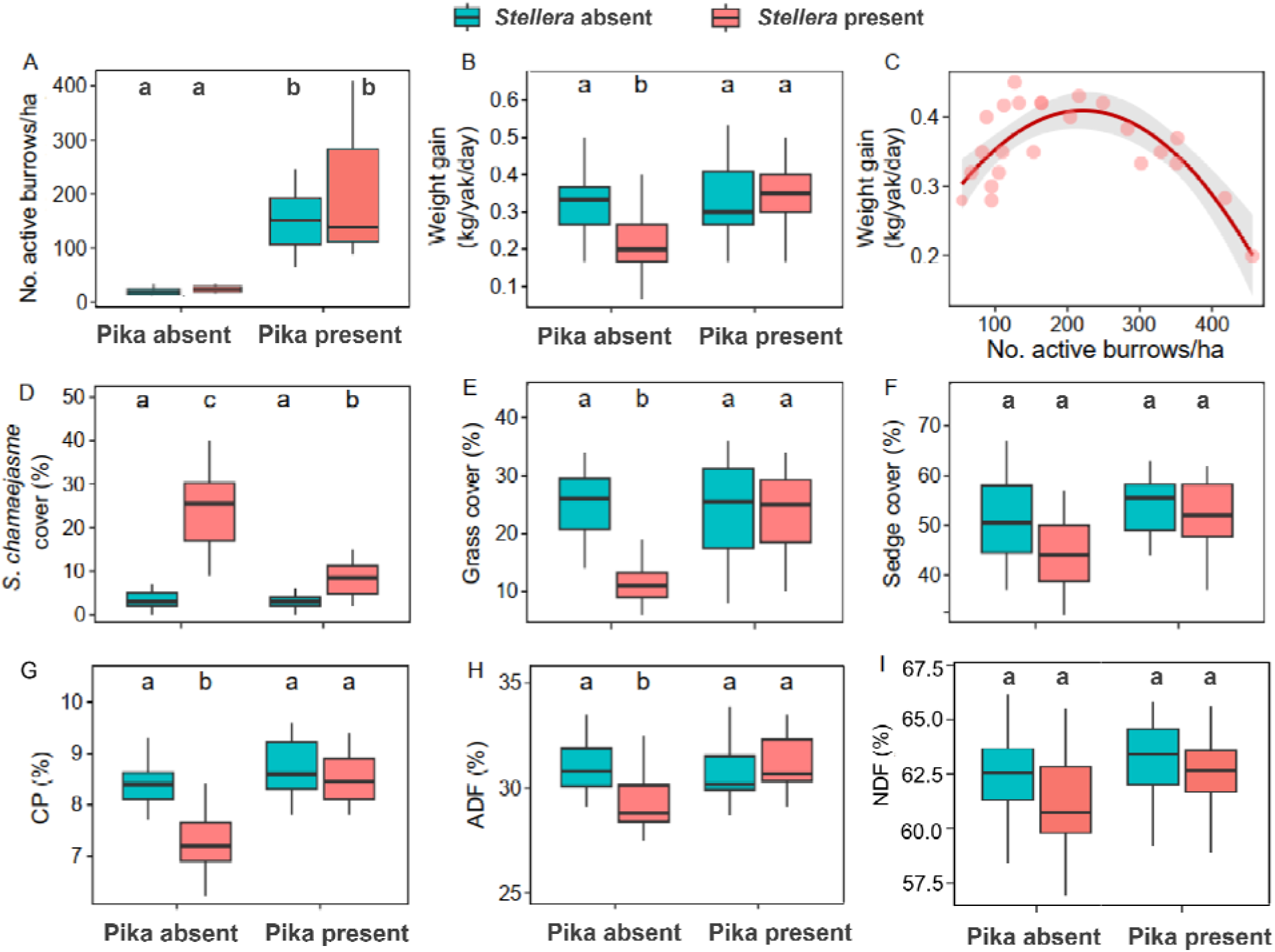
Combined effects of two-year (2022–2023) pika and *Stellera* removal treatments on yak growth, forage quantity, and quality in the field manipulative experiment. (**A**) Pika density (indicated by active burrow density), (**B**) yak weight gain, (**C**) relationship between pika density and yak weight gain, (**D**) *Stellera* cover, (**E**) grass cover, (**F**) sedge cover, (**G**) crude protein (CP) content, (**H**) acid detergent fiber (ADF) content, and (I) neutral detergent fiber (NDF) of total forage (dry mass basis). Average values from both years were used in analyses, yielding one data point per 150 × 150 m plot. Significant interactions between pikas and *Stellera* were evaluated using post hoc tests; means not sharing letters differ significantly. For panels (**A**) and (**F**), only main effects were significant (Table S5). Error bars indicate ± SE.

In the presence of *Stellera*, pika removal increased the cover abundance of this forb by two-fold (Fig. 3D, Table S5, S6), which led to decreases of 50% and 15% in cover abundances of co-occurring grasses and sedges, respectively (Fig. 3E,F, Table S5, S6). These shifts in vegetation composition also reduced the nutritional value of the major available forage for yaks, with decreases of 14% in crude protein (CP) content and 6% in acid detergent fiber (ADF) content in the pika-absent treatment compared with the pika-present treatment (Fig. 3G,H, Table S8, S9).

Pikas and *Stellera* had no interactive effects on abundance of sedges, forbs, and neutral detergent fiber (NDF) of total forage for yaks (Fig. 3F,I, Fig. S1, Table S5, S8). These results suggest that, in the presence of *Stellera*, pikas facilitate yak growth by suppressing poisonous plants, increasing both the availability and quality of palatable food plants for the livestock. Pika-yak facilitation was also linked to improved foraging efficiency in yaks when grazing alongside pikas. Bite rate (i.e., the number of bites taken on plants per hour) and bites/step ratio (i.e., the number of bites taken on plants per step) are two key indicators of foraging efficiency in large herbivores (Odadi et al., 2009). In the presence of *Stellera*, sedge bite rate and sedge bites per step of yaks significantly decreased by 32% and 46%, respectively (Fig. 4A,C, Table S10, S11), and grass bite rate and grass bites per step decreased similarly by 29% and 43% (Fig. 4B,D, Table S10, S11), in the pika-absent treatment compared to pika-present treatment. These decreases in yak foraging efficiency in the absence of pikas can be attributed to the increases in cover of the poisonous *Stellera* (Fig. 3D), which often act as a grazing deterrent and reduce access by yaks to palatable food items. Pikas and *Stellera* had no interactive effects on yaks’ foraging efficiency on forbs (Fig. S2, Table S10).

**Figure 4.**
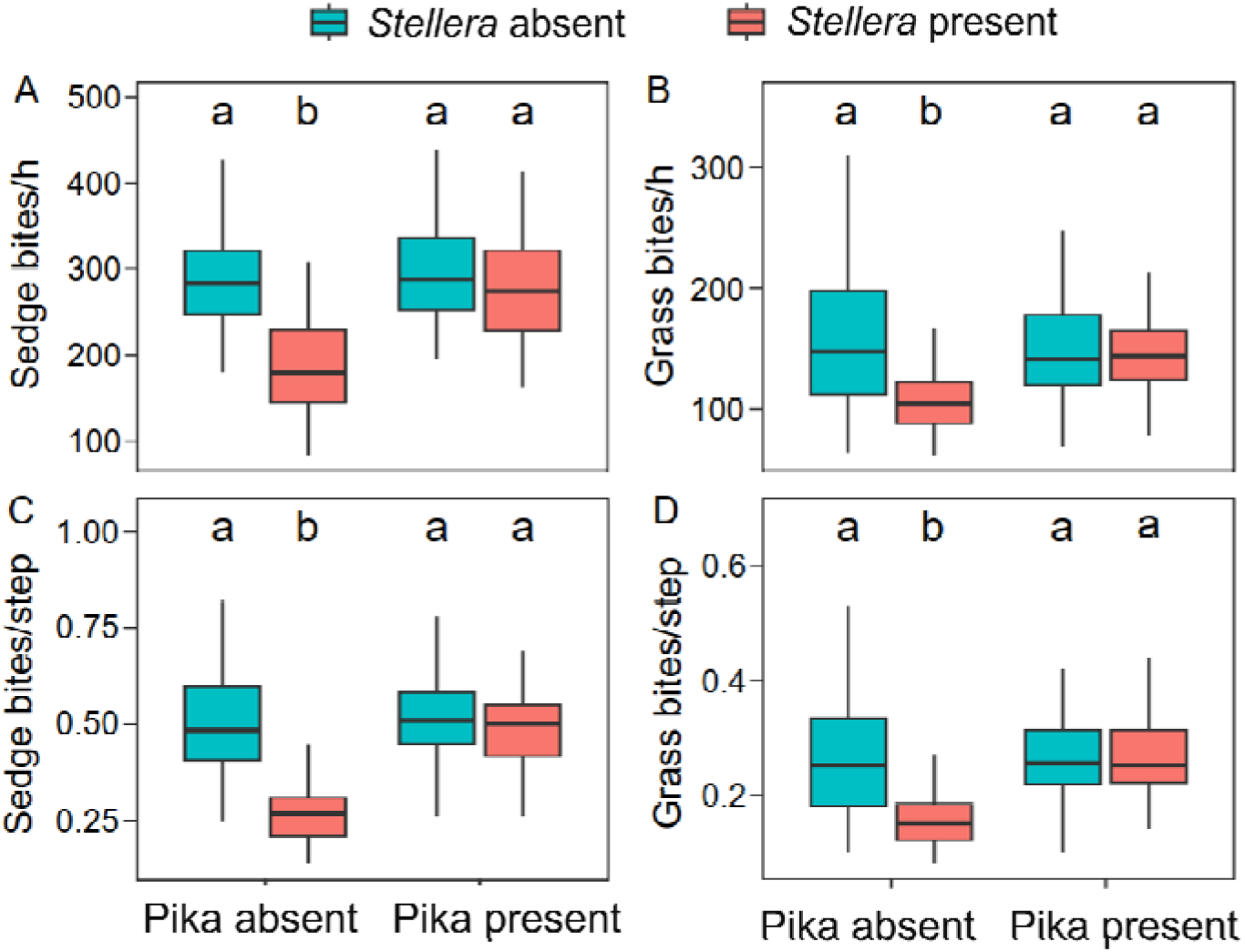
Combined effects of two-year (2022–2023) pika and *Stellera* removal treatments on yak foraging efficiency in the field manipulative experiment. (**A**, **B**) Bite rates and (**C**, **D**) bites per step for sedges and grasses, respectively. Mean values from both years were used for analysis (one data point per 150 × 150 m plot). Significant interactions between pikas and *Stellera* were evaluated using post hoc tests; means not sharing letters differ significantly. Error bars indicate ± SE.

## Discussion

By combining field surveys and manipulative experiments, we have demonstrated that pikas— when occurring at moderate densities—can benefit yak body growth by suppressing large poisonous plants and increasing the availability of palatable forage in the Qinghai-Tibetan Plateau. Our findings thus reconcile debates on the effects of pikas on yak production through changing vegetation properties. These observations provide new insights into how small and large mamamlian herbivores co-exist in a complex way in nature.

Ecosystem engineering is a key mechanism driving interspecific facilitation (Jones et al., 1994; Bruno et al., 2003; Machicote et al., 2004). Engineering activities of one herbivore species can indirectly benefit another by either increasing access to food resources or ameliorating abiotic conditions in particular habitats (Arsenault and Owen-Smith, 2002; Odadi et al., 2011; Zhong et al., 2014; Pan et al., 2019). In our system, the selective clipping activities of pikas greatly suppressed the abundance of large poisonous forbs (Fig. 3D), and increased both the quantity and quality of yaks’ palatable food plants (Fig. 3E-I). We speculate that these improvements in food availability and nutrition for yaks may be due to the release of grasses and sedges from competition with the poisonous forbs for limiting above- and below-ground resources (Lu et al., 2012; Li and Zhao, 2025). Pika-induced increases in food resources, along with fewer foraging barriers for yaks after poisonous forb removal by pikas, together increased foraging efficiency (Fig. 4) and thus facilitated weight gain by yaks (Fig. 3B). In addition to increasing food resources, the removal of poisonous plants by pikas may also benefit yaks by reducing the risk of incidental consumption (Lu et al., 2012; Li and Zhao, 2025), which can cause biochemical or physiological stresses in yaks that lead to sickness or even death (Keeler et al., 2013).

Our results add to a growing list of studies that highlight the importance of small mammal impacts on large herbivores. In grasslands of the North American Great Plains, for example, black-tailed prairie dogs (*Cynomys ludovicianus*) can alter growing-season forage quality and quantity both on and off their colonies, exerting either competitive or facilitatory impacts on daily forage intake rates and mass gain of cattle (O’Meilia et al., 1982; Detling 2006; Augustine and Derner, 2021; Augustine et al., 2024). By contrast, in drought years on the Mongolian mountain steppes, overlap in use of forage grasses (*Stipa krylovii* and *Agropyron cristatum*) can lead to competition for food between Mongolian pika (*Ochotona pallasi*) and goats, sheep, and cattle, with potential negative impacts on livestock production (Retzer, 2007). Given the ubiquity of small mammals, and their ability to strongly modify plant and soil properties via herbivory and ecosystem engineering, the impacts of small mammals on co-occurring large herbivores may be complex, resulting in either positive or negative outcomes which would benefit from further investigations.

Our results reveal the importance of wild small herbivores in counteracting livestock grazing-induced vegetation imbalance in rangelands. The coexistence of a diverse of herivore species with different diet selections and size classes can lead to an “compensatory effect” on grass and forb biomass that helps to maintain a balance and diverse plant community (Ritchie and Olff, 1999), with importance consequences for ecosystem functioning and services. In our system, the widespread of *Stellera* across the Qinghai-Tibetan Plateau is itself a product of long-term grazing pressure, which suppresses palatable grasses and sedges while releasing unpalatable, stress-tolerant forbs (Lu et al., 2012; Li and Zhao, 2025). The pika’s selective clipping of forbs helps promoting graminoid recovery, benefiting not only livestock production but may also other key rangeland functions (e.g., carbon storage and nutrient cycling) (Lu et al., 2012; Li et al., 2023; Li and Zhao, 2025). Such modification effects of small herbivores, especially those colonial and hyper-abundant ones (e.g., rodents and insects), on large grazers’ ecological functions may be more common and importance than previously perceived (Davidson et al., 2010, 2012; Goheen et al., 2010; Clark et al., 2012; Zhong et al., 2014).

Because of the natural variations in pika density in the pika-present treatment, we were able to obtain a hump-shaped relationship between yak weight gains and pika burrow densities in these plots. Compared with the absence of pikas, the facilitative effect reached its maximum at approximately 200 burrows/ha but became competitive at densities exceeding 400 burrows/ha (Fig. 3C). This result reveals that pika density modulates the net outcome for yak weight gain, with a facilitation peak at ∼200 burrows/ha and a competition onset above 400 burrows/ha. Our findings offer empirical evidence for the non-monotonicity theory, under which the competition-facilitation balance varies with population density: facilitation dominates at low densities, competition at high densities, and these density-dependent shifts may underpin community stability and productivity (Zhang, 2003; Zhang et al., 2015). The theory further holds that the facilitation threshold, not the competition-facilitation transition, is the critical factor governing the stability of interacting species or communities (Zhang et al., 2015).

Collectively, our experiments provide empirical evidence that small mammals can facilitate large herbivores through altering vegetation properties. By suppressing tall poisonous forbs, plateau pikas improved forage composition and quality, increased yak foraging efficiency, and ultimately boosting yak weight gain. These findings reconcile the debate that small mammalian herbivores act as rangeland pests in outbreaking years and as beneficial ecological engineers in non-outbreaking years (Fan et al., 1998, 1999; Zhang and Wang, 1998; Singleton et al., 1999, 2010; Smith and Foggin, 1999; Delibes-Mateos et al., 2011). Our study demonstrates that small mammalian herbivores enhance pastoral systems at moderate densities but impair them at high densities. Thus, at burrow densities below 400 burrows/ha, no pika population management is required in the study region.

Despite of these, several questions remain deserve further investigation. First, our study examined pika–yak interactions only during the summer period, when food resources are most abundant. Whether such facilitative effects weaken or even shift toward competition under more stressful conditions—for example, when forage becomes limited during autumn or winter— remains to be tested. Second, if the documented facilitation of yaks by pikas prompts herders to increase yak densities, could the resulting rise in livestock herbivory push pika populations beyond the levels observed here, potentially toward the threshold where facilitation gives way to competition (Yang et al., 2026)? Third, our study site located at approximately 3,200 m elevation, was relatively low by Qinghai-Tibetan Plateau standards. *Stellera* becomes less common at elevations > 4,000 m, where a majority of livestock grazing occurs. It would be instructive to learn, through follow-up studies, whether similar facilitation occurs where unpalatable (and mildly poisonous) species in such genera as *Astragalus*, *Oxytropis*, and *Thermopsis* replace *Stellera* as the problematic plants for pastoralists (Lu et al., 2012; Li and Zhao, 2025). Finally, it is unclear whether similar facilitation as observed here applies to the other principal livestock species in the area, such as domestic sheep and goats.

Current rangeland management practices often rely heavily on rodenticides and other toxic compounds, leading to widespread small-mammal eradication (Fan et al., 1998, 1999; Zhang and Wang, 1998; Singleton et al., 1999, 2010). Our findings show that coexistence when small herbivorous mammals at low to moderate densities can enhance livestock growth performance. Recognizing and managing such facilitatory interactions between herbivore guilds supports international goals to integrate biodiversity conservation with food security and climate adaptation. Rangeland policy should therefore move from pest eradication toward ecologically based management (Singleton et al., 1999) that regulates herbivore populations through ecosystem processes and habitat management. Such an approach embraces the full spectrum of herbivores—from rodents to megafauna to livestock—as contributors to multifunctional landscapes in the Anthropocene (cf. Svenning et al., 2024).

## Methods

All pika and yak manipulations were carried out in accordance with the Law of the People’ s Republic of China on the Protection of Wildlife (1988).

### Study system and background

We conducted the study at a grazing grassland in Menyuan County, Qinghai Province, China (37°48’ N, 101°56’ E, 3200 m a.s.l.). The site is located in the northeast Qinghai-Tibetan Plateau and has continental cold/humid climate conditions, with a summer rainy season and a winter dry season. The mean annual temperature is 4.2°C, and rainfall was 750 mm. The vegetation is typical of alpine meadows. The grassland is dominated by sedges such as *Kobresia* spp. (e.g., *K. humilis* and *K. graminifolia*), subdominant species include grasses such as *Elymus* spp. (i.e., *E. nutans*), *Festuca* spp. (e.g., *F. ovina*), and *Stipa* spp. (i.e., *S. aliena*), and companion species of forbs such as *Potentilla* spp. (e.g., *P. anserina* and *P. multifida*) and *Medicago ruthenica*. In recent decades, the poisonous *Stellera chamaejasme* forb, commonly named “wolf poison”, which is toxic to livestock, has encroached and has become a dominant plant owing to human disturbances and climate changes, competing with forage plants for shared resources such as light, soil water, and nutrients (Lu et al., 2012; Li and Zhao, 2025).

The plateau pika (*Ochotona curzoniae*) is a small (body length ca. 120-190 mm, and body weight ca. 110-170 g) lagomorph endemic to and dominant in the alpine meadows of the Qinghai-Tibetan Plateau (Qu et al., 2013). When populations outbreak, pika population densities can reach over 500-1000 active burrows per hectare (Fan et al., 1998,1999; Zhang et al., 1999). However, our study grassland hosts a moderate density (ca. 100-300 active burrows per hectare) of plateau pikas during the forage growing seasons (June to August), providing us with an ideal opportunity to investigate the effects of non-outbreak pika density on yak performance. Pikas live in social groups, breed during the warm summer and have lifespans about 120-250 days (Qu et al., 2013). The plateau pika is a keystone species due to the feeding, forb-clipping, and burrowing activities of individuals that exert profound effects on soils, plants, and other animals in the meadows (Smith et al., 2019). By selectively feeding on and clipping large dicotyledonous plants such as forbs (Jiang and Xia, 1985; Liu et al., 2008), pikas occupy a different plant-food utilization niche compared with domestic livestock such as yaks and sheep, which often prefer monocotyledonous plants such as grasses and sedges (Pan et al., 2019; Li et al., 2023). Notably, pikas often clip (although do not eat) the poisonous *S. chamaejasmehas* because its relatively large size hinders predator detection (Fan et al., 1998). This species of pika prefer habitats where vegetation height is low than where it is higher. Wild ungulates, such as Tibetan gazelles (*Procapra picticaudata*) (Harris et al., 2015), and other small mammals such as rabbits and zokors occur rarely in the area.

Tibetan yak (*Bos grunniens*) is the major ruminant species on the Tibetan rangelands due to the species’ excellent adaptability and production performance (Long et al., 2008). It is estimated that about 14 million domestic yaks live on the Tibetan Plateau, providing local herdsmen with daily necessities like meat, milk, wool, skins, fuel and economic benefit (Dong et al., 2020). In our study site, the grassland has been managed by pastoralism of domesticated yaks for years, and the grazing intensity is mainly controlled by pastoral practice.

### Field survey #1: Diet preferences of pikas and yaks

In July (peak growing season) 2021, we investigated the diet/clipping selection of pikas and yaks in the study site. For pika feeding/clipping preferences, we randomly selected 10 2 m × 2 m plots separated by at least 100 m from each other in the study site. A large cage enclosure 1.5 m high and 2 m × 2 m bottom surface area, covered with a 5 × 5 mm plastic mesh window screen, was assigned to each plot. Before cage installation, we carefully checked the plants within each plot and removed those that had been previously grazed or damaged by herbivores. We then captured pikas nearby and placed one adult into each enclosure, allowing each animal to freely feed/clip plants within the cage for 20 min before it was released. We then laid out a 2 m linear transect in each cage plot consisting of 10 0.2 × 0.2 m quadrats. If vegetation had been consumed (i.e., plant tissues were removed and digested) or clipped (i.e., plant tissues were cut down and lay on the ground surface), we assigned that quadrat a value of one for that vegetation group (i.e., *Stellera*, sedges, grasses, and forbs); if there was no sign of consumption, a value of zero was assigned. Values assigned for each vegetation group were summed for the transect and divided by 10 to obtain a frequency of feeding/clipping use ranging from 0% to 100% (Clark et al., 2012). To document yak diet composition, we placed 10 1 × 1 m quadrats on the ground at approximately 20 m intervals along 10 250 m transects that were randomly located on fresh grazing paths of yaks. We recorded and calculated how frequently different plant groups were grazed by yaks using the same methods as above.

### Field survey #2: Associations among pikas, poisonous plants, and yak activities

In July 2021, we investigated the potential facilitation of pikas on yaks mediated by the poisonous *Stellera* forbs under unmanipulated field conditions in the study site. We firstly selected 30 10 m × 10 m plots separated by at least 200 m from each other in the study site. The site was grazed by yaks at low to moderate intensities (i.e., 0.5-1.5 animal units/ha), with varying abundance of pikas and the poisonous *Stellera* forbs. Within each plot, we then assessed the abundance of pikas and *Stellera*, and foraging activities of yaks. We visually counted the number of active burrows (hole entrances characterized by clear openings, fresh soil or pika feces) to indicate pika abundance. For poisonous plant abundance, we visually estimated the percentage of the ground surface covered by *Stellera* in four 1 m × 1 m quadrats within each plot. For yaks, we recorded the number of dungs present in each plot, which is regarded a good measure for assaying grazing pressure in grasslands (Eldridge et al., 2025).

### Field manipulative experiment: Interactive effects of pika and poisonous plants on yak performance

#### Experimental design

In May 2022, we established four replicate blocks of experimental plots, for a total of 16 plots with similar plant community composition and pika densities in the study site (Table S12). The site had not been disturbed by human activities (e.g., grazing or mowing) for two years prior to the initiation of the study. Each block had the following 2 × 2 factorial design: presence of pikas and poisonous plants, pikas only, poisonous plants only, and where neither pikas nor poisonous plants was present. Plot treatments were randomly assigned within each block. Minimum distances between the four replicate blocks of plots were 200–300 m. Each of the four plots in a replicate block was separated by 50 m, and each plot was 150 × 150 m. To avoid edge effects, we sampled plant, pika, and yak variables within the 100 × 100 m area at the center of each plot.

At the start of experimental treatments each year, we obtained 32 yak steers aged 2 years and weighing 115 ± 7.8 kg from adjacent and/or nearby pastures, and randomly grouped them into 16 herds of two yaks each. We then randomly allocated these yak herds to the 16 experimental plots (one herd/plot), creating a light to moderate grazing intensity recommended by local government guidelines, which often allow yak to maximize their growth rates (Dong et al., 2003). Electric fencing was used to confine yaks within the experimental plots. To align with their local grazing habits, yaks were allowed to graze daily in the experimental plots between 08:00 and 18:00, after which the animals were removed from the plots and housed in shelters overnight without feeding. All yaks had free access to fresh water and a mineral-lick block (Cangzhou Leysin Biotechnology Co., Ltd, Cangzhou, China) during the experimental period. Yak grazing under this regime continued for two growing seasons (June to August) in 2022 and 2023.

#### Pika treatments

We installed exclosures to control pika populations in the plots. Initially, pikas were present on all plots, and the treatments with pika absence were implemented by removing pikas and preventing recolonization by fencing using an iron sheet around the perimeter of the plots. The iron sheet extended 0.60 m aboveground to prevent pikas jumping in or out, and was buried 1.5 m below the soil surface to deter animals from burrowing underneath.

In May 2022, after the establishment of the iron sheet fences, the pika exclusion treatments were initiated by removing pikas from the allocated experimental plots. For the no-pika treatment, pikas were trapped once every two weeks using 30 live traps (25 cm high × 25 cm wide × 40 cm long) within each plot and relocated elsewhere in the study site. Pika activities (e.g., number of active burrows) in the experimental plots were monitored monthly (June to August) thereafter, and pikas entering the exclosure plots were removed as necessary to maintain the exclusion treatments.

#### Poisonous plant treatments

For the experimental plots without poisonous plants, we clipped and removed *Stellera* forbs using garden clippers. To simulate the clipping behavior of pikas, we clipped only those *Stellera* forbs exceeding 20 cm in height. This threshold was chosen based on previous observations that pikas preferentially target large forbs of this size (Liu et al., 2009). We removed these poisonous plants once a month from June to August. For the experimental plots with poisonous plants, *Stellera* forbs were left intact.

#### Yak body growth and grazing activity

During the growing seasons (June to August) in 2022 and 2023, we recorded the initial and final body weights (Weighbridge, Shanghai Jiujin Electronics Apparatus Co. Shanghai, China) of yaks each month to calculate their average daily weight gain. Each month, we also observed the grazing activities of each yak for six 2-hour focal periods. During these observations, we recorded the number of bites that the yaks took on different forage plant groups (i.e., sedges, grasses, and forbs) and the number of steps they took. The foraging efficiency of yaks on each plant group (bites/step) was calculated as the number of bites on specific plant group/the total number of steps during the observations (Odadi et al., 2009).

#### Forage quantity and quality

From June to August, along with the measurements of yak behavior and attributes as described above, we assessed how the food resources of yaks changed in the experimental plots each month. To assess forage quantity, we randomly assigned ten 1 × 1 m quadrats spaced at least 10 m from each other, and recorded the percentage of the ground surface covered by each plant group (i.e., *Stellera*, sedges, grasses, and forbs). To assess forage quality, five forage samples were collected from each grazing plot to quantify their nutritive values. To obtain samples that reflect the forage actually consumed by yaks, we tracked the animals along their grazing paths and collected the plant tissues of the two most frequently consumed species: the dominant sedge *Kobresia humilis* and the dominant grass *Elymus nutans* (Fig. 2B; Pan et al., 2019). The collected tissues of each species were dried in a forced-air oven at 60‘ ° C for 48‘ h, then ground through a 1-mm mesh. Subsequently, 5‘ g of each dried and ground species were combined in a 1:1 dry mass ratio, and the resulting mixture was stored in plastic bags for subsequent analyses.

We analysed forage samples for crude protein (CP), acid detergent fiber (ADF), and neutral detergent fiber (NDF). Total nitrogen content was determined by the Kjeldahl method (2300; Foss Tecator AB, Hoganas, Sweden) using selenium as the catalyst, and CP was calculated as 6.25 × nitrogen. ADF and NDF were analysed with an automatic fibertec apparatus (M6, Foss Tecator AB, Hoganas, Sweden) by the method of Van Soest et al. (1991).

### Statistical analyses

All data were analyzed using linear models, generalized linear mixed models, or generalized additive models, with the choice of model and statistical family guided by the structure and distribution of the data. Post-hoc comparisons were conducted only when the pika × *Stellera* interaction term was significant. For the 2021 field surveys, we fitted generalized linear mixed models with plot and month as random effects. We then used generalized additive mixed models for the cover abundance of *Stellera* and active pika burrow density, with plot as a random effect, and linear regression models for dung density and *Stellera* cover. For the field manipulation experiments in 2022 and 2023, we constructed generalized linear mixed models with the dependent variables regressed against the interactive effect of pika and *Stellera* treatments, while including block, year, and month as random effects to capture the hierarchical structure of the data. Models assumed Gaussian, beta (for proportions), or Tweedie (for non-normal data) distributions, selected based on data type and model fit. There were two models that used the tweedie family (forb and sedge bite rate). We chose to use tweedie because a normal gaussian family model fitted the results poorly. A significance threshold of *P* = 0.05 was applied, with Tukey-HSD or Sidak post-hoc tests used where appropriate. All data management, modeling, and visualization approaches were carried out in R, with dependencies managed using *renv*. The main modeling packages were *glmmTMB* (Brooks et al., 2017) and *mgcv* (Wood, 2017), with *DHARMa* (Hartig, 2022) used for model diagnostics. Data management relied on the *tidyverse* (Wickham et al., 2019) suite of packages. A summary of all statistical models used in the study is available in Table S13. A complete record of package versions is available in the renv.lock file in the repository Zenodo: https://doi.org/10.5281/zenodo.18290921.

## Funding

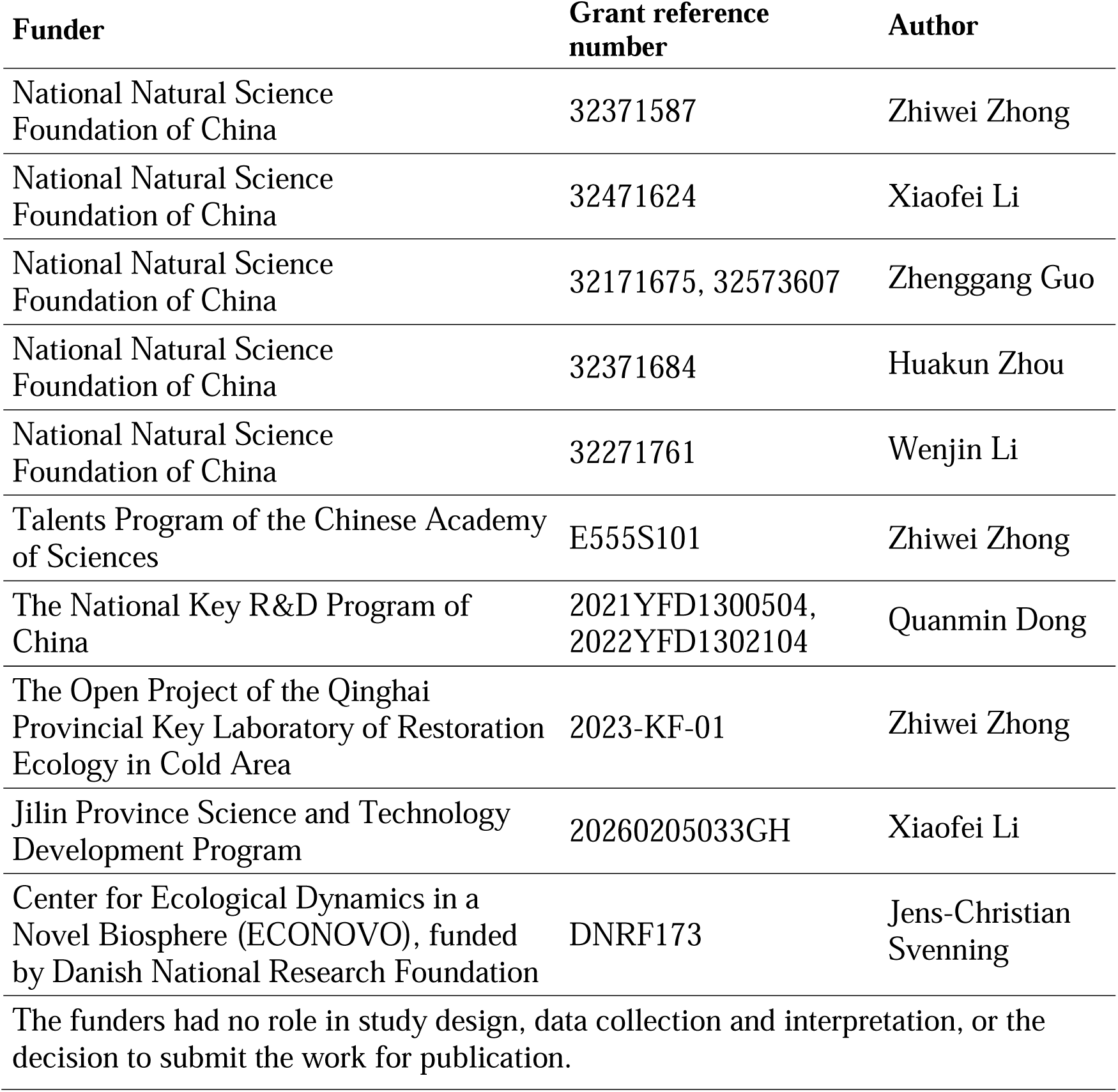

## Author contributions

Zhiwei Zhong, Conceptualization, Investigation, Visualization, Methodology, Funding acquisition, Writing-original draft, Writing-review and editing; Bingbo Ni, Data curation, Investigation; Douglas Lawton, Software, Formal analysis, Writing-original draft, Writing-review and editing; Xiaofei Li, Data curation, Investigation, Funding acquisition; Xiaona Zheng, Data curation, Investigation; Huakun Zhou, Writing-review and editing; Junhu Su, Writing-review and editing; Wenjin Li, Writing-review and editing; Fujiang Hou, Writing-review and editing; Zhenggang Guo, Writing-review and editing; Quanmin Dong, Writing-review and editing; Shikui Dong, Writing-review and editing; Christopher R. Dickman, Writing-review and editing; Jens-Christian Svenning, Conceptualization, Supervision, Writing-original draft, Writing-review and editing; Ying Gao, Conceptualization, Investigation, Resources, Writing-review and editing; Zhibin Zhang, Conceptualization, Supervision, Writing-review and editing

## Data availability

All data are available in ddlawton/pika_yak_interactions: initial release for peer review, version v1.0.0, Zenodo, https://doi.org/10.5281/zenodo.18290921, GitHub link: https://github.com/ddlawton/pika_yak_interactions

## Supplementary Materials for

Materials and Methods

Figs. S1, S2

Tables S1 to S13

**Figure S1.**
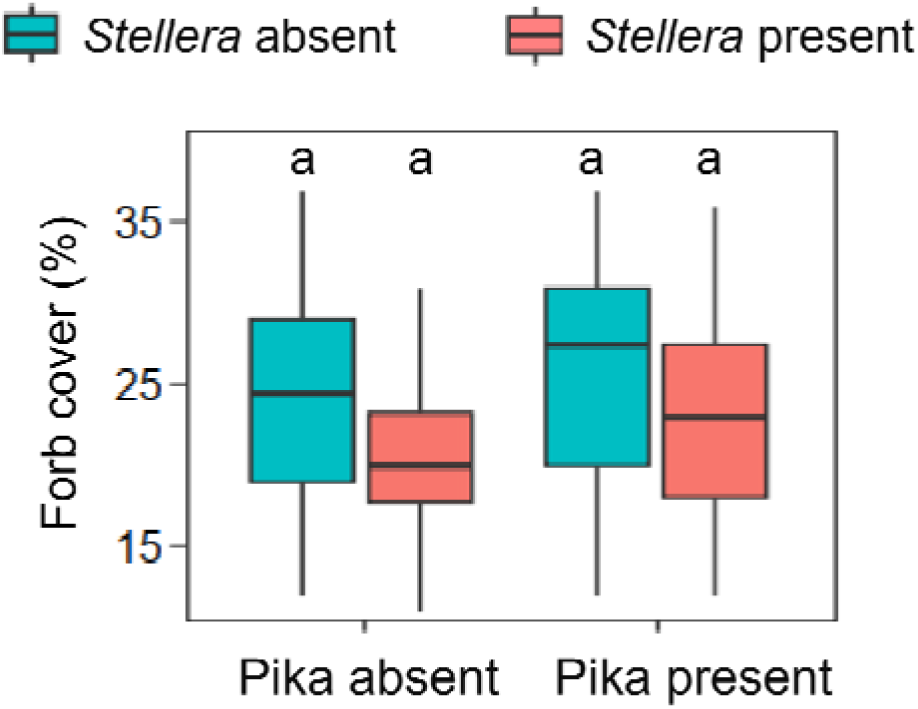
Combined effects of two-year (2022–2023) pika and *Stellera* removal on forb cover in the field manipulative experiment. The average values of each variable in the two years were used for statistical analysis, providing a single data point for each variable in each 150 × 150 m plot. Error bars represent ± SE.

**Figure S2.**
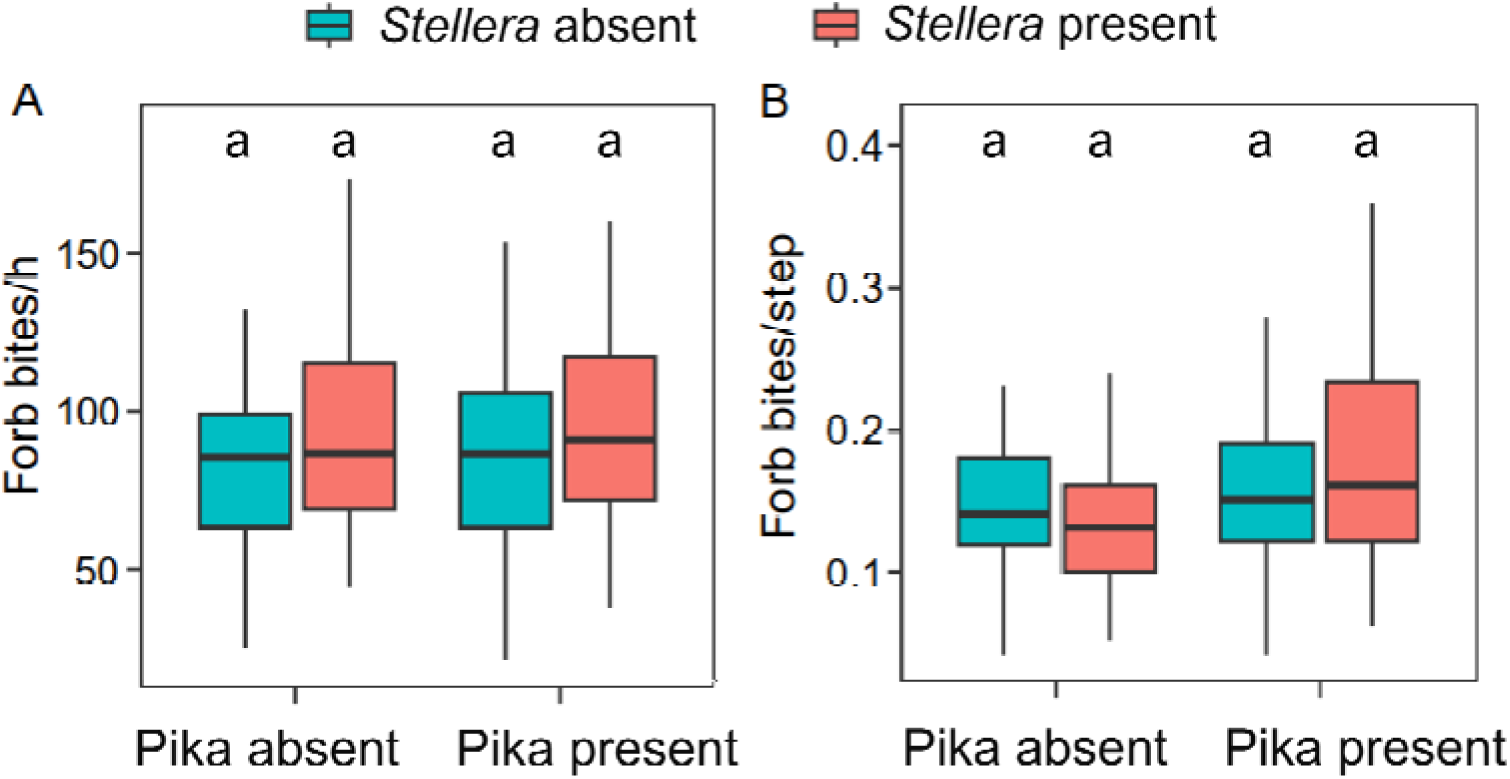
Combined effects of two-year (2022–2023) pika and *Stellera* removal on foraging efficiency of yaks in the field manipulative experiment. (**A**) bite rates and (**B**) bites per step of yaks for forbs, respectively. The average values of each variable in the two years were used for statistical analysis, providing a single data point for each variable in each 150 × 150 m plot. Error bars represent ± SE.

**Table S1.**
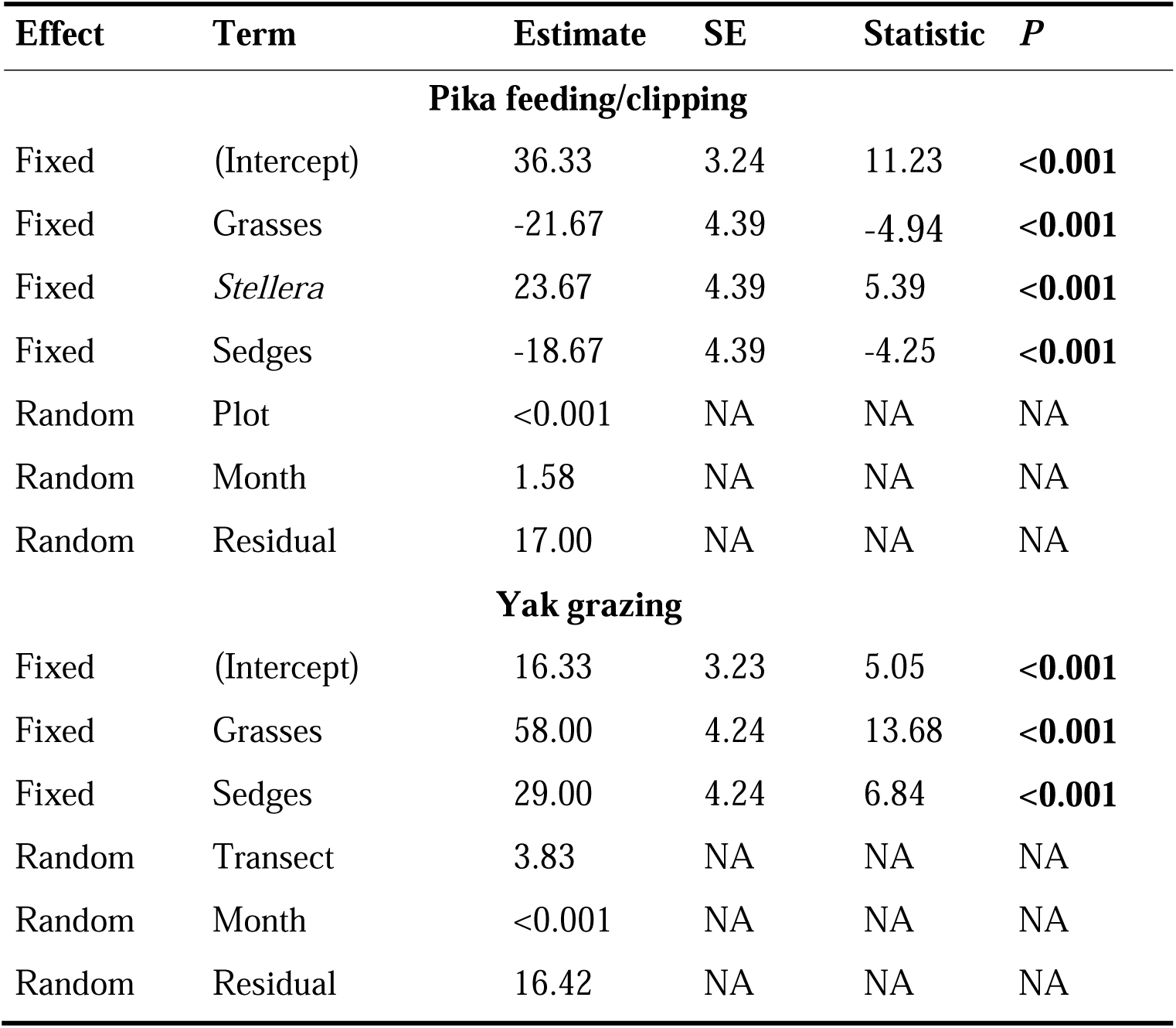
Model summary for diet and clipping selections of pikas and yaks in the field surveys in July 2021.

**Table S2.** Model contrasts for diet and clipping selections of pikas and yaks in the field surveys.

| Contrast | Estimate | SE | df | *t* ratio | *P* |
| --- | --- | --- | --- | --- | --- |
| **Pika feeding/clipping** |  |  |  |  |  |
| Forbs–Grasses | 21.67 | 4.39 | 113.00 | 4.94 | <**0.001** |
| Forbs–*Stellera* | -23.67 | 4.39 | 113.00 | -5.39 | <**0.001** |
| Forbs–Sedges | 18.67 | 4.39 | 113.00 | 4.25 | <**0.001** |
| Grasses–*Stellera* | -45.33 | 4.39 | 113.00 | -10.33 | <**0.001** |
| Grasses–Sedges | -3.00 | 4.39 | 113.00 | -0.68 | 0.90 |
| *Stellera*–Sedges | 42.33 | 4.39 | 113.00 | 9.64 | <**0.001** |
| **Yak grazing** |  |  |  |  |  |
| Forbs–Grasses | -58.00 | 4.24 | 84.00 | -13.68 | <**0.001** |
| Forbs–Sedges | -29.00 | 4.24 | 84.00 | -6.84 | <**0.001** |
| Grasses–Sedges | 29.00 | 4.24 | 84.00 | 6.84 | <**0.001** |

**Table S3.** Model summary for *Stellera* cover regressed on active pika burrow density. Estimates are from a generalized additive mixed model (Gaussian family) with plot as a random effect (n = 30 plots). density.

| Term | Estimate | SE | Statistic | *P* | edf | ref.df |
| --- | --- | --- | --- | --- | --- | --- |
| Intercept | 13.33 | 1.21 | 10.99 | < **0.00** | NA | NA |
| Active pika burrow density | NA | NA | 13.17 | < **0.00** | 2.10 | 7.00 |
| Plot | NA | NA | 1.17 | 0.06 | 15.12 | 29.00 |

**Table S4.**
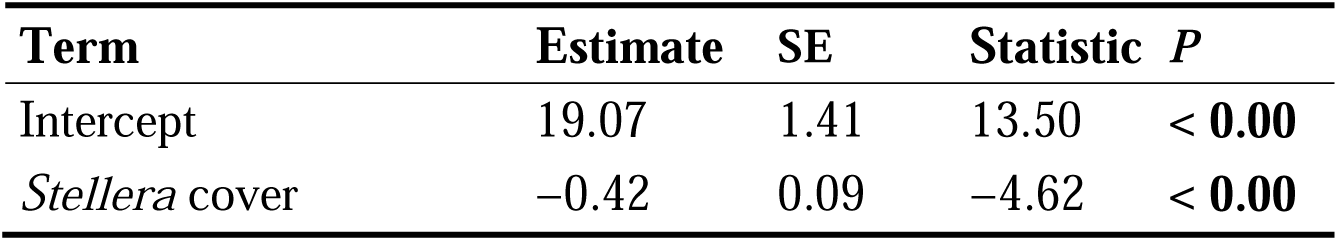
Model summary for yak dung density regressed on *Stellera* cover. Estimates are from a linear regression model (n = 30 plots).

| **Term** | **Estimate** | **SE** | **Statistic** | **P** |
| --- | --- | --- | --- | --- |
| Intercept | 19.07 | 1.41 | 13.50 | < **0.00** |
| *Stellera* cover | −0.42 | 0.09 | −4.62 | < **0.00** |

**Table S5.**
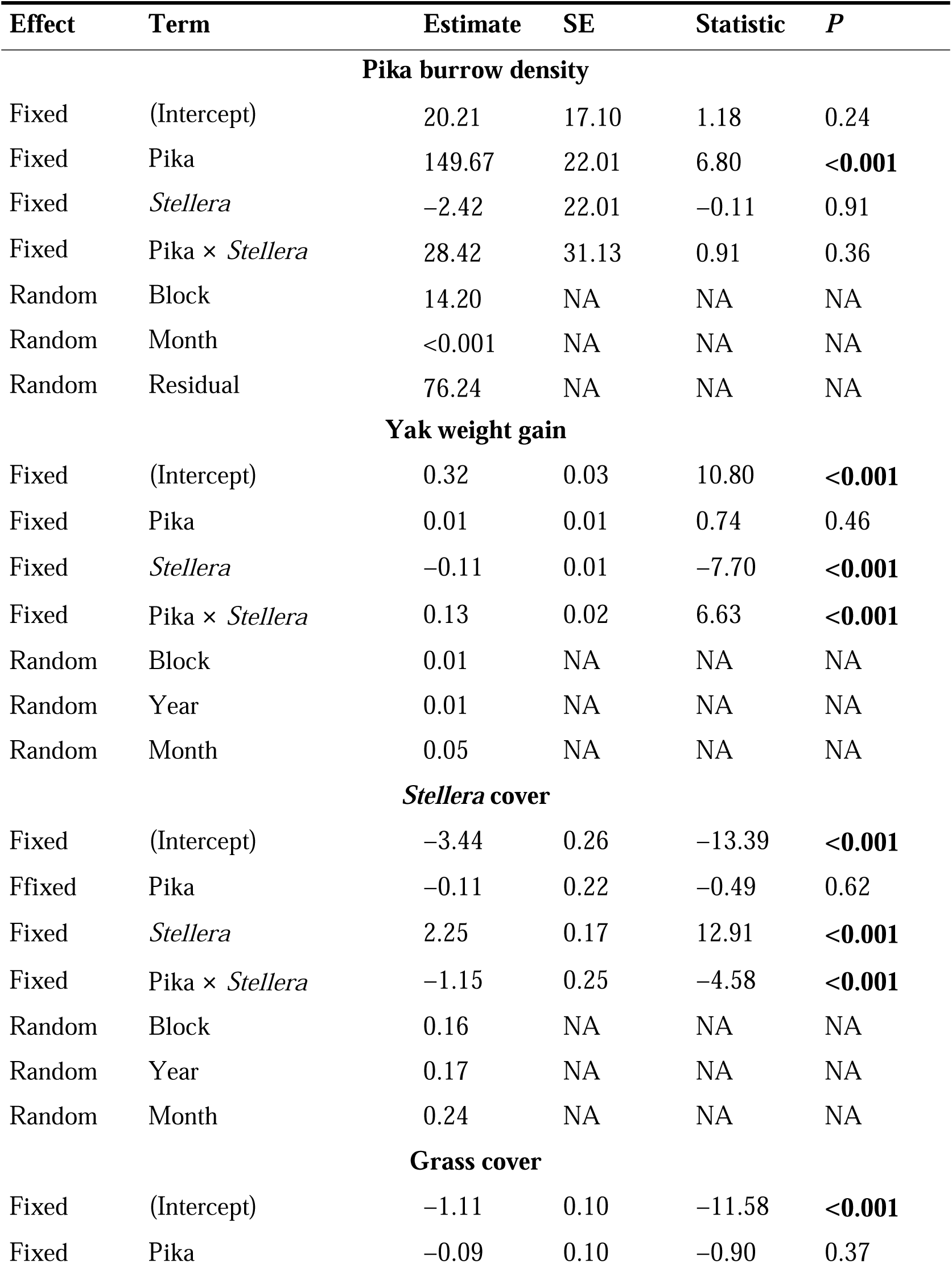
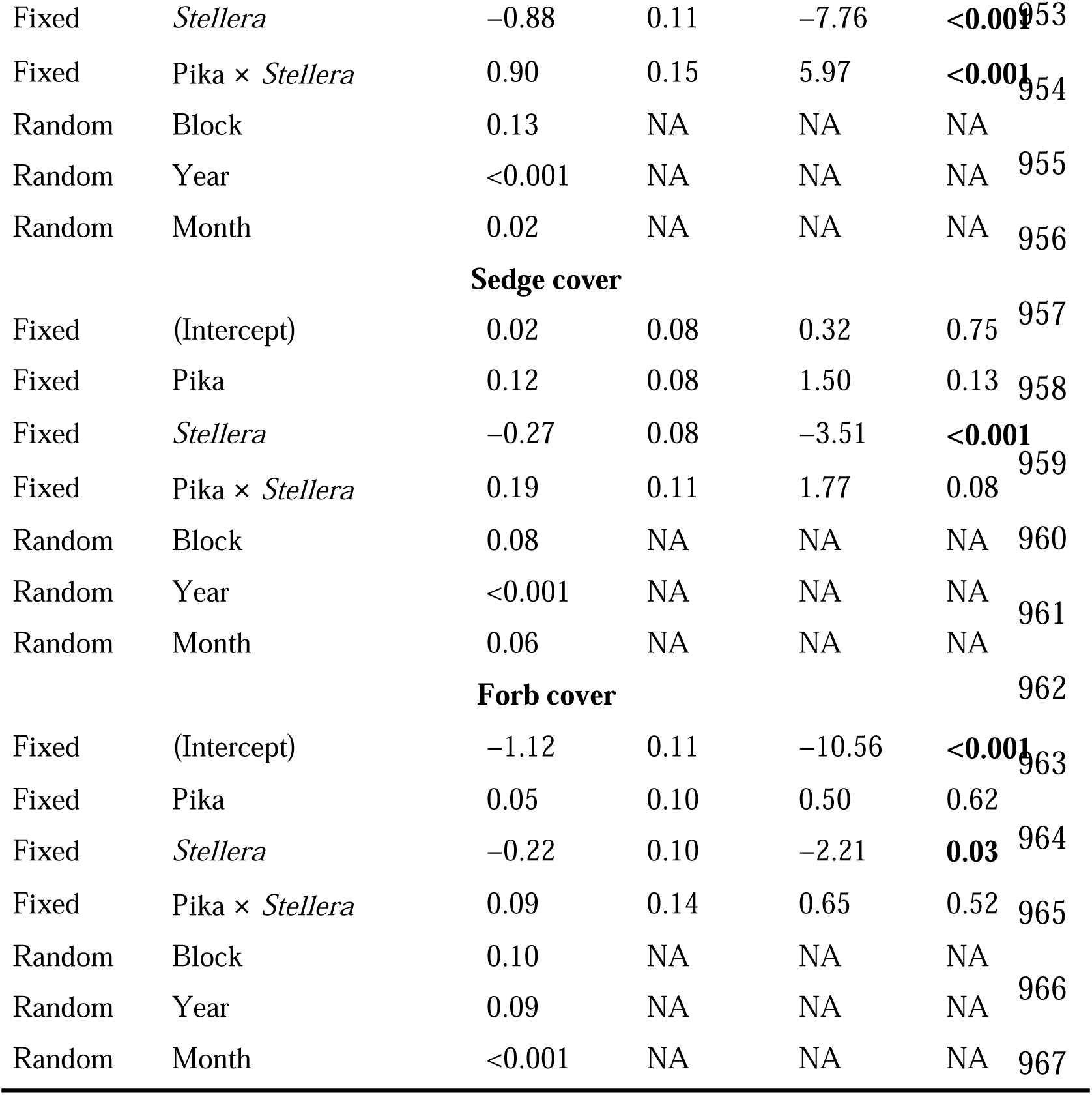
Model summary for interactive effects of pikas and *Stellera* on pika burrow density, yak weight gain, *Stellera* cover, grass cover, sedge cover, and forb cover in the field manipulative experiment in 2022-2023.

**Table S6.**
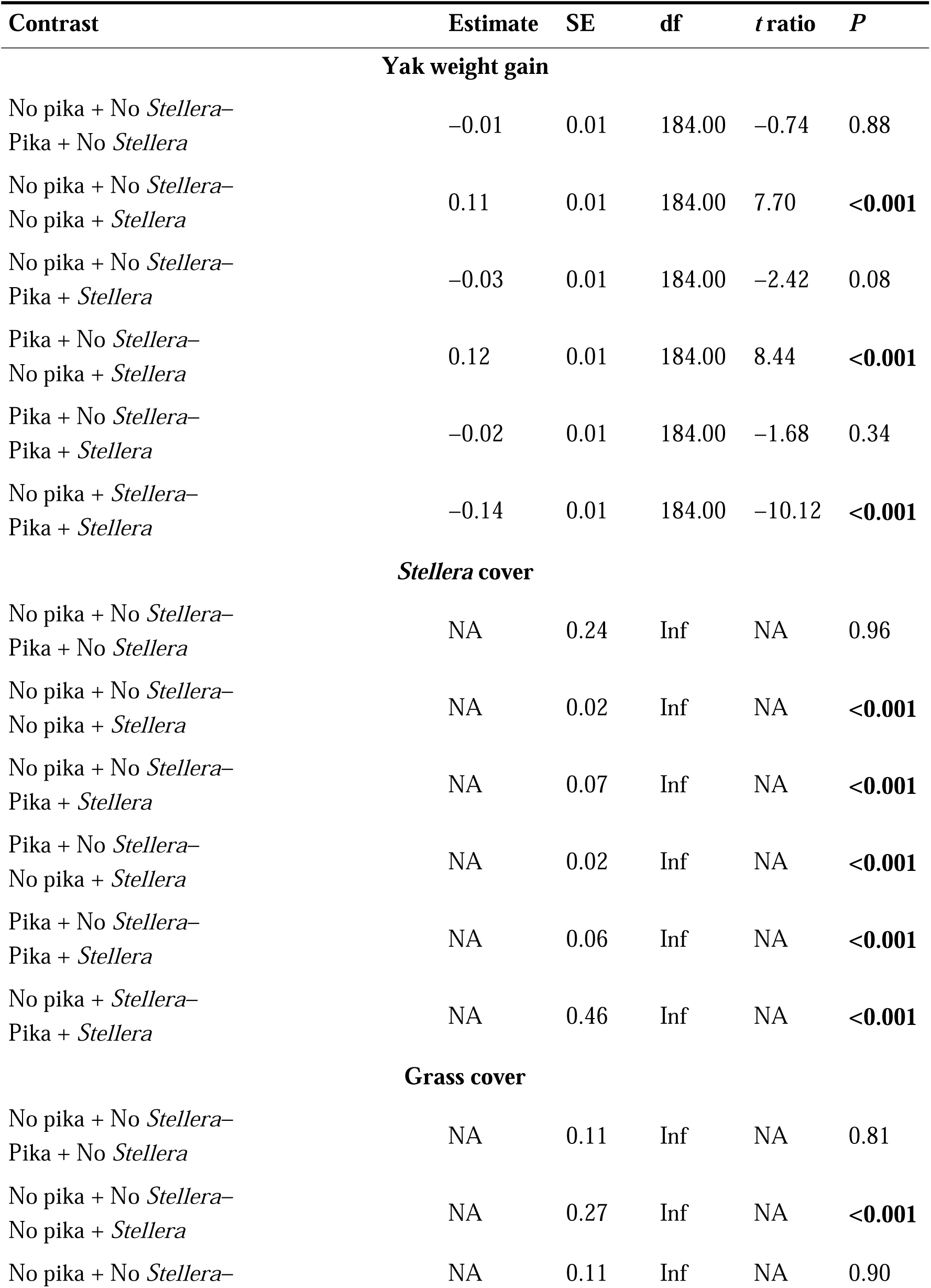
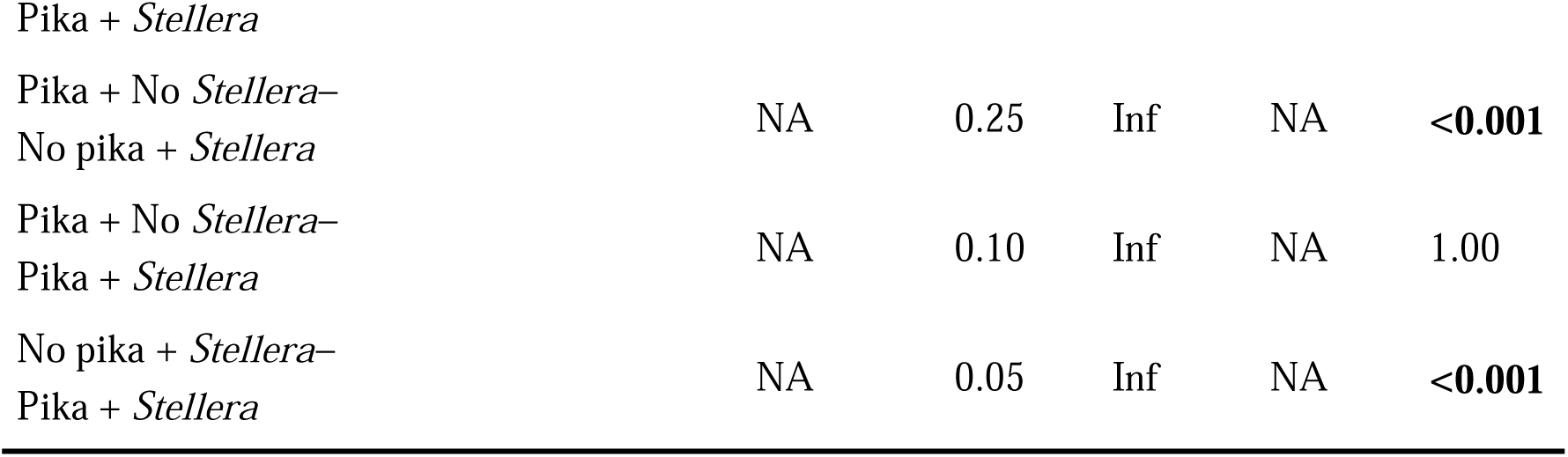
Model contrasts for yak weight gain, *Stellera* cover, and grass cover in the field manipulative experiment in 2022-2023.

**Table S7.** Model summary for yak weight gain regressed on active burrow count in plots with both pika presence and *Stellera* presence. Estimates are from a generalized additive mixed model (Gaussian family) with a non-linear smooth for active burrows, block, year, and month as random effects (n = 24 observations from the pika + *Stellera* treatment).

| Term | edf | ref.df | statistic | P |
| --- | --- | --- | --- | --- |
| Active burrows | 2.27 | 14.00 | 4.74 | <0.001 |
| Block | 1.72 | 3.00 | 1.44 | 0.09 |
| Year | 0.23 | 1.00 | 0.31 | 0.25 |
| Month | 1.57 | 2.00 | 5.24 | 0.01 |

**Table S8.** Model summary for interactive effects of pikas and *Stellera* on crude protein (CP) %, acid detergent fiber (ADF) %, and neutral detergent fiber (NDF) % of total forage in the field manipulative experiment in 2022-2023.

| Effect | Term | Estimate | SE | Statistic | P |
| --- | --- | --- | --- | --- | --- |
| **CP** |  |  |  |  |  |
| Fixed | (Intercept) | -2.39 | 0.02 | -107.03 | **<0.001** |
| Fixed | Pika | 0.03 | 0.02 | 1.88 | 0.06 |
| Fixed | *Stellera* | -0.15 | 0.02 | -8.72 | **<0.001** |
| Fixed | Pika × *Stellera* | 0.13 | 0.02 | 5.45 | **<0.001** |
| Random | Block | 0.01 | NA | NA | NA |
| Random | Year | <0.001 | NA | NA | NA |
| Random | Month | 0.03 | NA | NA | NA |
| **ADF** |  |  |  |  |  |
| Fixed | (Intercept) | -0.82 | 0.03 | -32.36 | **<0.001** |
| Fixed | Pika | 0.01 | 0.02 | 0.46 | 0.65 |
| Fixed | *Stellera* | -0.06 | 0.02 | -3.14 | **<0.001** |
| Fixed | Pika × *Stellera* | 0.08 | 0.03 | 2.84 | **<0.001** |
| Random | Block | 0.03 | NA | NA | NA |
| Random | Year | <0.001 | NA | NA | NA |
| Random | Month | 0.03 | NA | NA | NA |
| **NDF** |  |  |  |  |  |
| Fixed | (Intercept) | 0.51 | 0.02 | 27.44 | **<0.001** |
| Fixed | Pika | 0.03 | 0.03 | 1.03 | 0.30 |
| Fixed | *Stellera* | -0.05 | 0.02 | -2.03 | **0.04** |
| Fixed | Pika × *Stellera* | 0.02 | 0.04 | 0.61 | 0.54 |
| Random | Block | 0.01 | NA | NA | NA |
| Random | Year | <0.001 | NA | NA | NA |
| Random | Month | <0.001 | NA | NA | NA |

**Table S9.** Model contrasts for crude protein (CP) % and acid detergent fiber (ADF) % of total forage in the field manipulative experiment in 2022-2023.

| Contrast | Estimate | SE | df | t ratio | P |
| --- | --- | --- | --- | --- | --- |
| CP |  |  |  |  |  |
| No pika + No *Stellera*–<br>Pika + No *Stellera* | NA | 0.02 | Inf | NA | 0.24 |
| No pika + No *Stellera*–<br>No pika + *Stellera* | NA | 0.02 | Inf | NA | <0.001 |
| No pika + No *Stellera*–<br>Pika + *Stellera* | NA | 0.02 | Inf | NA | 0.91 |
| Pika + No *Stellera*–<br>No pika + *Stellera* | NA | 0.02 | Inf | NA | <0.001 |
| Pika + No *Stellera*–<br>Pika + *Stellera* | NA | 0.02 | Inf | NA | 0.61 |
| No pika + *Stellera*–<br>Pika + *Stellera* | NA | 0.01 | Inf | NA | <0.001 |
| ADF |  |  |  |  |  |
| No pika + No *Stellera*–<br>Pika + No *Stellera* | NA | 0.02 | Inf | NA | 0.97 |
| No pika + No *Stellera*–<br>No pika + *Stellera* | NA | 0.02 | Inf | NA | <0.001 |
| No pika + No *Stellera*–<br>Pika + *Stellera* | NA | 0.02 | Inf | NA | 0.54 |
| Pika + No *Stellera*–<br>No pika + *Stellera* | NA | 0.02 | Inf | NA | <0.001 |
| Pika + No *Stellera*–<br>Pika + *Stellera* | NA | 0.02 | Inf | NA | 0.82 |
| No pika + *Stellera*–<br>Pika + *Stellera* | NA | 0.02 | Inf | NA | <0.001 |

**Table S10.** Model summary for interactive effects of pikas and *Stellera* on yak bite rate (bites/h) and the bite to step ratio (bites/step) for grasses, sedges, and forbs in the field manipulative experiment in 2022-2023.

| Effect | Term | Estimate | SE | Statistic | P |
| --- | --- | --- | --- | --- | --- |
| **Grass bites/h** |  |  |  |  |  |
| Fixed | (Intercept) | 151.72 | 7.49 | 20.25 | **<0.001** |
| Fixed | Pika | -0.80 | 8.61 | -0.09 | 0.93 |
| Fixed | *Stellera* | -45.34 | 8.61 | -5.27 | **<0.001** |
| Fixed | Pika × *Stellera* | 43.95 | 12.17 | 3.61 | **<0.001** |
| Random | Block | 4.78 | NA | NA | NA |
| Random | Year | 5.17 | NA | NA | NA |
| Random | Month | 0.00 | NA | NA | NA |
| Random | Residual | 42.16 | NA | NA | NA |
| **Sedge bites/h** |  |  |  |  |  |
| Fixed | (Intercept) | 5.65 | 0.04 | 138.78 | **<0.001** |
| Fixed | Pika | 0.04 | 0.04 | 0.92 | 0.36 |
| Fixed | *Stellera* | -0.42 | 0.05 | -8.71 | **<0.001** |
| Fixed | Pika × *Stellera* | 0.35 | 0.06 | 5.51 | **<0.001** |
| Random | Block | 0.04 | NA | NA | NA |
| Random | Year | 0.02 | NA | NA | NA |
| Random | Month | 0.03 | NA | NA | NA |
| **Forb bites/h** |  |  |  |  |  |
| Fixed | (Intercept) | 4.39 | 0.07 | 65.12 | **<0.001** |
| Fixed | Pika | 0.07 | 0.07 | 1.06 | 0.29 |
| Fixed | *Stellera* | 0.13 | 0.07 | 1.90 | 0.06 |
| Fixed | Pika × *Stellera* | -0.04 | 0.10 | -0.44 | 0.66 |
| Random | Block | 0.06 | NA | NA | NA |
| Random | Year | 0.00 | NA | NA | NA |
| Random | Month | 0.05 | NA | NA | NA |
| **Grass bites/step** |  |  |  |  |  |
| Fixed | (Intercept) | -1.01 | 0.08 | -13.16 | **<0.001** |
| Fixed | Pika | 0.02 | 0.09 | 0.19 | 0.85 |
| Fixed | *Stellera* | -0.60 | 0.10 | -6.28 | **<0.001** |
| Fixed | Pika × *Stellera* | 0.67 | 0.13 | 5.15 | <0.001 |
| Random | Block | 0.03 | NA | NA | NA |
| Random | Year | 0.06 | NA | NA | NA |
| Random | Month | 0.00 | NA | NA | NA |
| **Sedge bites/step** |  |  |  |  |  |
| Fixed | (Intercept) | 0.04 | 0.14 | 0.28 | 0.78 |
| Fixed | Pika | 0.08 | 0.12 | 0.67 | 0.50 |
| Fixed | *Stellera* | −0.95 | 0.12 | −7.61 | <0.001 |
| Fixed | Pika × *Stellera* | 1.01 | 0.17 | 5.83 | <0.001 |
| Random | Block | 0.17 | NA | NA | NA |
| Random | Year | 0.08 | NA | NA | NA |
| Random | Month | 0.06 | NA | NA | NA |
| **Forb bites/step** |  |  |  |  |  |
| Fixed | (Intercept) | −1.76 | 0.09 | −18.74 | <0.001 |
| Fixed | Pika | 0.06 | 0.09 | 0.64 | 0.52 |
| Fixed | *Stellera* | −0.05 | 0.09 | −0.49 | 0.63 |
| Fixed | Pika × *Stellera* | 0.24 | 0.13 | 1.83 | 0.07 |
| Random | Block | 0.11 | NA | NA | NA |
| Random | Year | 0.00 | NA | NA | NA |
| Random | Month | 0.07 | NA | NA | NA |

**Table S11.**
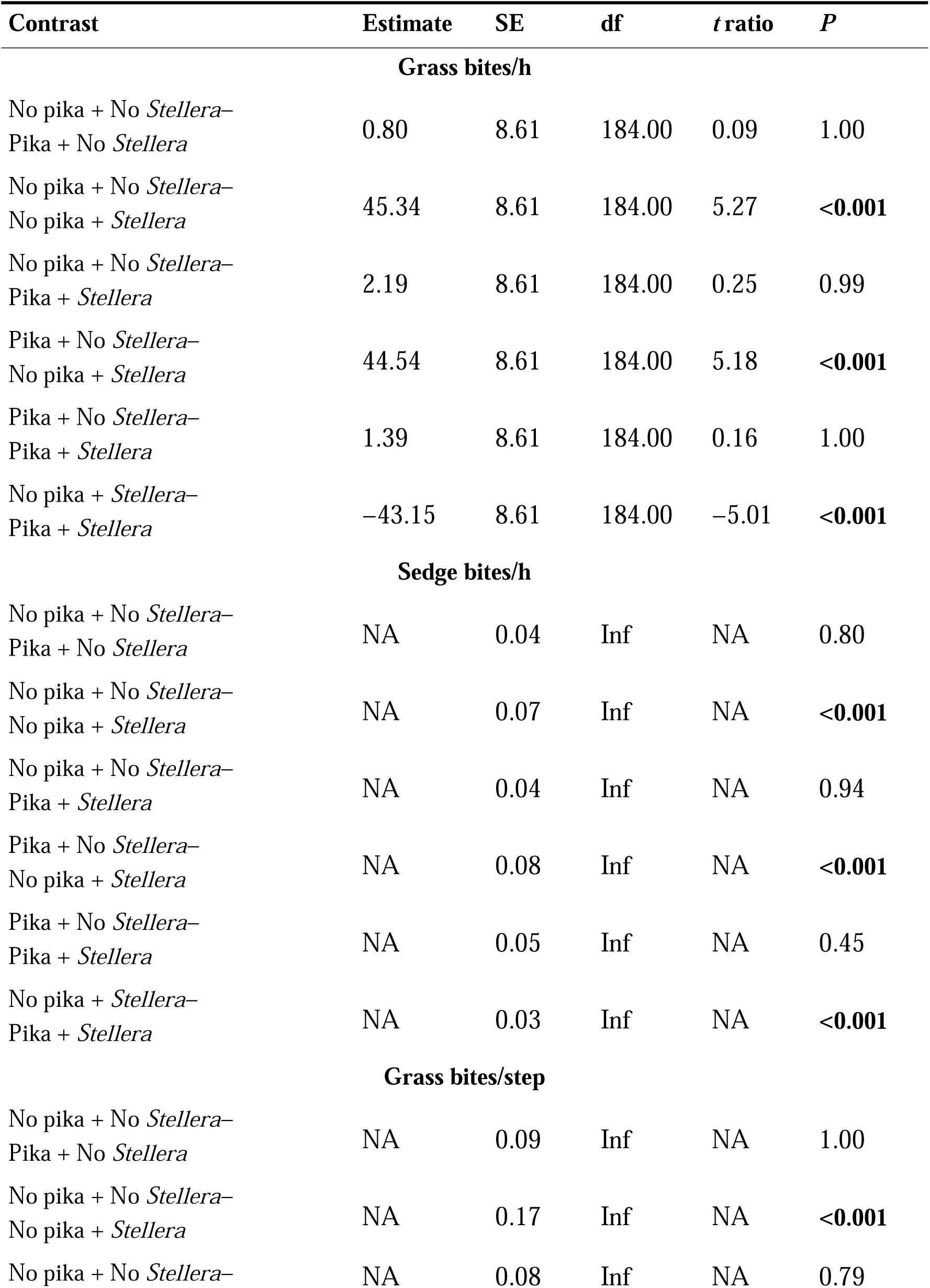
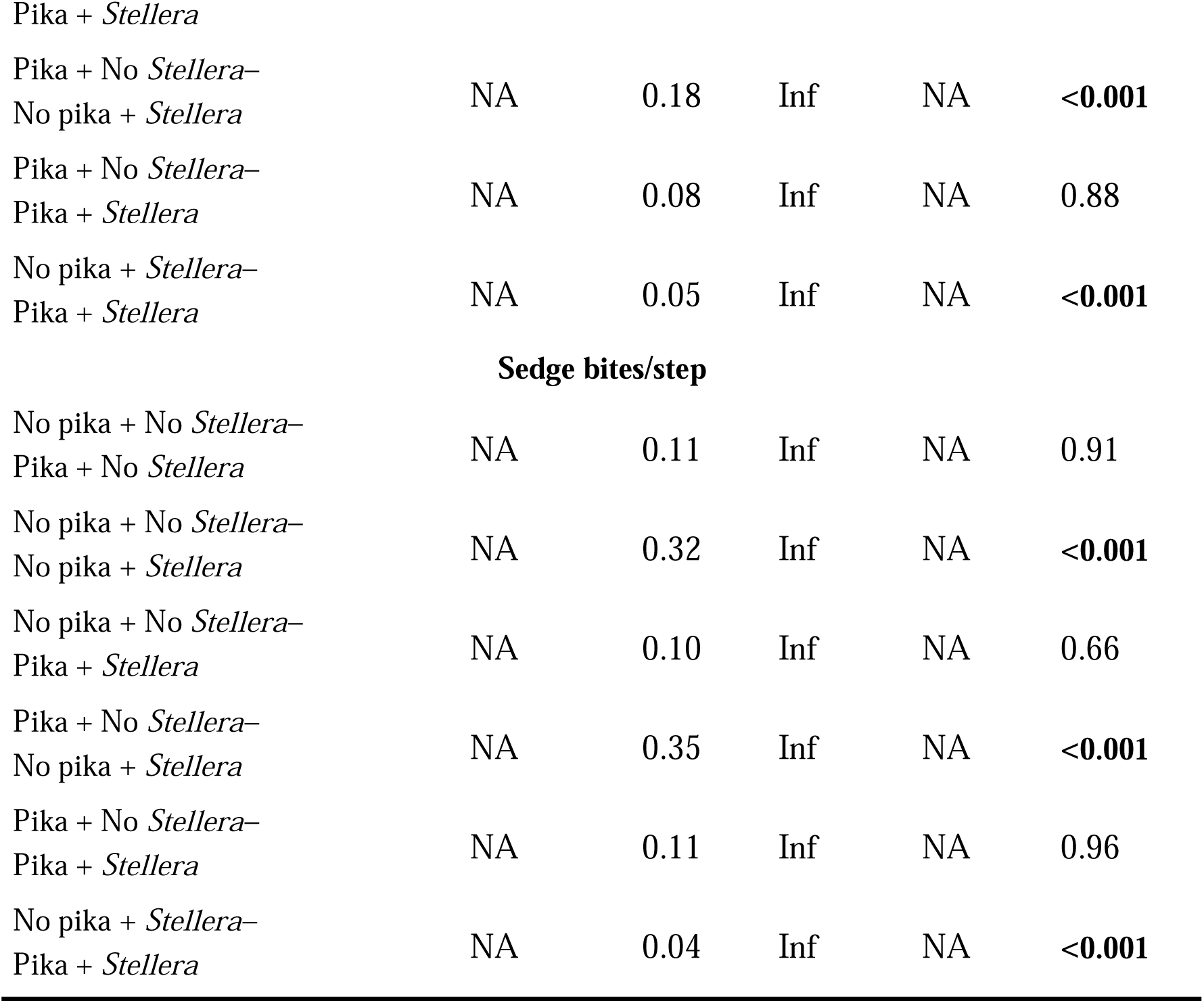
Model contrasts for yak bite rate (bites/h) and the bite to step ratio (bites/step) on grasses and sedges in the field manipulative experiment in 2022-2023.

| Contrast | Estimate | SE | df | *t* ratio | *P* |
| --- | --- | --- | --- | --- | --- |
| **Grass bites/h** |  |  |  |  |  |
| No pika + No *Stellera*–<br>Pika + No *Stellera* | 0.80 | 8.61 | 184.00 | 0.09 | 1.00 |
| No pika + No *Stellera*–<br>No pika + *Stellera* | 45.34 | 8.61 | 184.00 | 5.27 | <0.001 |
| No pika + No *Stellera*–<br>Pika + *Stellera* | 2.19 | 8.61 | 184.00 | 0.25 | 0.99 |
| Pika + No *Stellera*–<br>No pika + *Stellera* | 44.54 | 8.61 | 184.00 | 5.18 | <0.001 |
| Pika + No *Stellera*–<br>Pika + *Stellera* | 1.39 | 8.61 | 184.00 | 0.16 | 1.00 |
| No pika + *Stellera*–<br>Pika + *Stellera* | –43.15 | 8.61 | 184.00 | –5.01 | <0.001 |
| **Sedge bites/h** |  |  |  |  |  |
| No pika + No *Stellera*–<br>Pika + No *Stellera* | NA | 0.04 | Inf | NA | 0.80 |
| No pika + No *Stellera*–<br>No pika + *Stellera* | NA | 0.07 | Inf | NA | <0.001 |
| No pika + No *Stellera*–<br>Pika + *Stellera* | NA | 0.04 | Inf | NA | 0.94 |
| Pika + No *Stellera*–<br>No pika + *Stellera* | NA | 0.08 | Inf | NA | <0.001 |
| Pika + No *Stellera*–<br>Pika + *Stellera* | NA | 0.05 | Inf | NA | 0.45 |
| No pika + *Stellera*–<br>Pika + *Stellera* | NA | 0.03 | Inf | NA | <0.001 |
| **Grass bites/step** |  |  |  |  |  |
| No pika + No *Stellera*–<br>Pika + No *Stellera* | NA | 0.09 | Inf | NA | 1.00 |
| No pika + No *Stellera*–<br>No pika + *Stellera* | NA | 0.17 | Inf | NA | <0.001 |
| No pika + No *Stellera*– | NA | 0.08 | Inf | NA | 0.79 |
| Pika + No *Stellera*–<br>No pika + *Stellera* | NA | 0.18 | Inf | NA | <**0.001** |
| Pika + No *Stellera*–<br>Pika + *Stellera* | NA | 0.08 | Inf | NA | 0.88 |
| No pika + *Stellera*–<br>Pika + *Stellera* | NA | 0.05 | Inf | NA | <**0.001** |
| **Sedge bites/step** |  |  |  |  |  |
| No pika + No *Stellera*–<br>Pika + No *Stellera* | NA | 0.11 | Inf | NA | 0.91 |
| No pika + No *Stellera*–<br>No pika + *Stellera* | NA | 0.32 | Inf | NA | <**0.001** |
| No pika + No *Stellera*–<br>Pika + *Stellera* | NA | 0.10 | Inf | NA | 0.66 |
| Pika + No *Stellera*–<br>No pika + *Stellera* | NA | 0.35 | Inf | NA | <**0.001** |
| Pika + No *Stellera*–<br>Pika + *Stellera* | NA | 0.11 | Inf | NA | 0.96 |
| No pika + *Stellera*–<br>Pika + *Stellera* | NA | 0.04 | Inf | NA | <**0.001** |

**Table S12.**
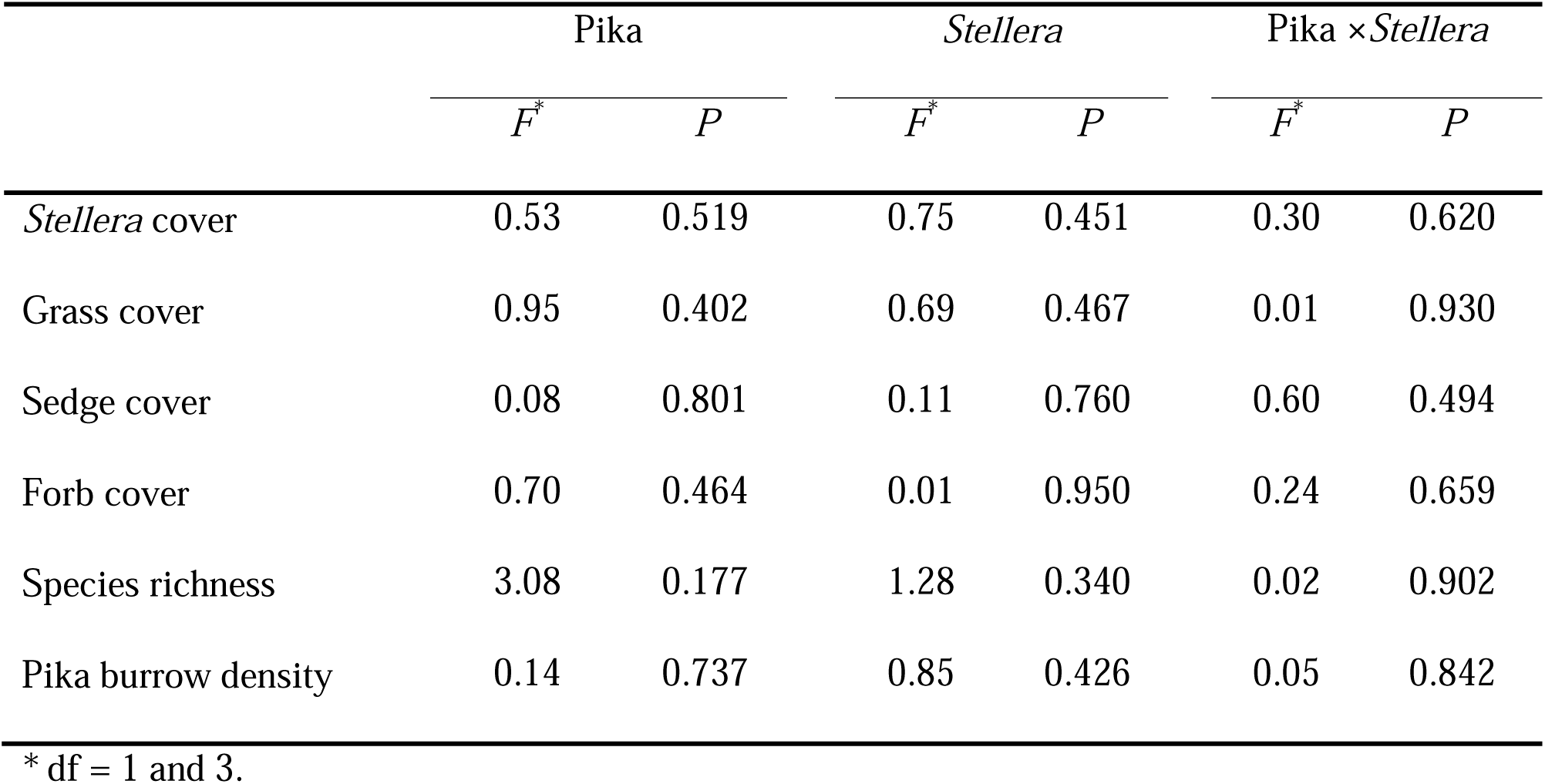
Summary statistics from generalized linear mixed effects models testing for pre-treatment differences in response variables representing plant community composition (cover by group) and diversity in designated mesocosm locations, as measured in 2021 prior to initiation of pika presence/absence and *Stellera* presence/absence treatments.

|  | Pika |  | *Stellera* |  | Pika × *Stellera* |  |
| --- | --- | --- | --- | --- | --- | --- |
|  | *F*<sup>*</sup> | *P* | *F*<sup>*</sup> | *P* | *F*<sup>*</sup> | *P* |
| *Stellera* cover | 0.53 | 0.519 | 0.75 | 0.451 | 0.30 | 0.620 |
| Grass cover | 0.95 | 0.402 | 0.69 | 0.467 | 0.01 | 0.930 |
| Sedge cover | 0.08 | 0.801 | 0.11 | 0.760 | 0.60 | 0.494 |
| Forb cover | 0.70 | 0.464 | 0.01 | 0.950 | 0.24 | 0.659 |
| Species richness | 3.08 | 0.177 | 1.28 | 0.340 | 0.02 | 0.902 |
| Pika burrow density | 0.14 | 0.737 | 0.85 | 0.426 | 0.05 | 0.842 |

\* df = 1 and 3.

**Table S13.**
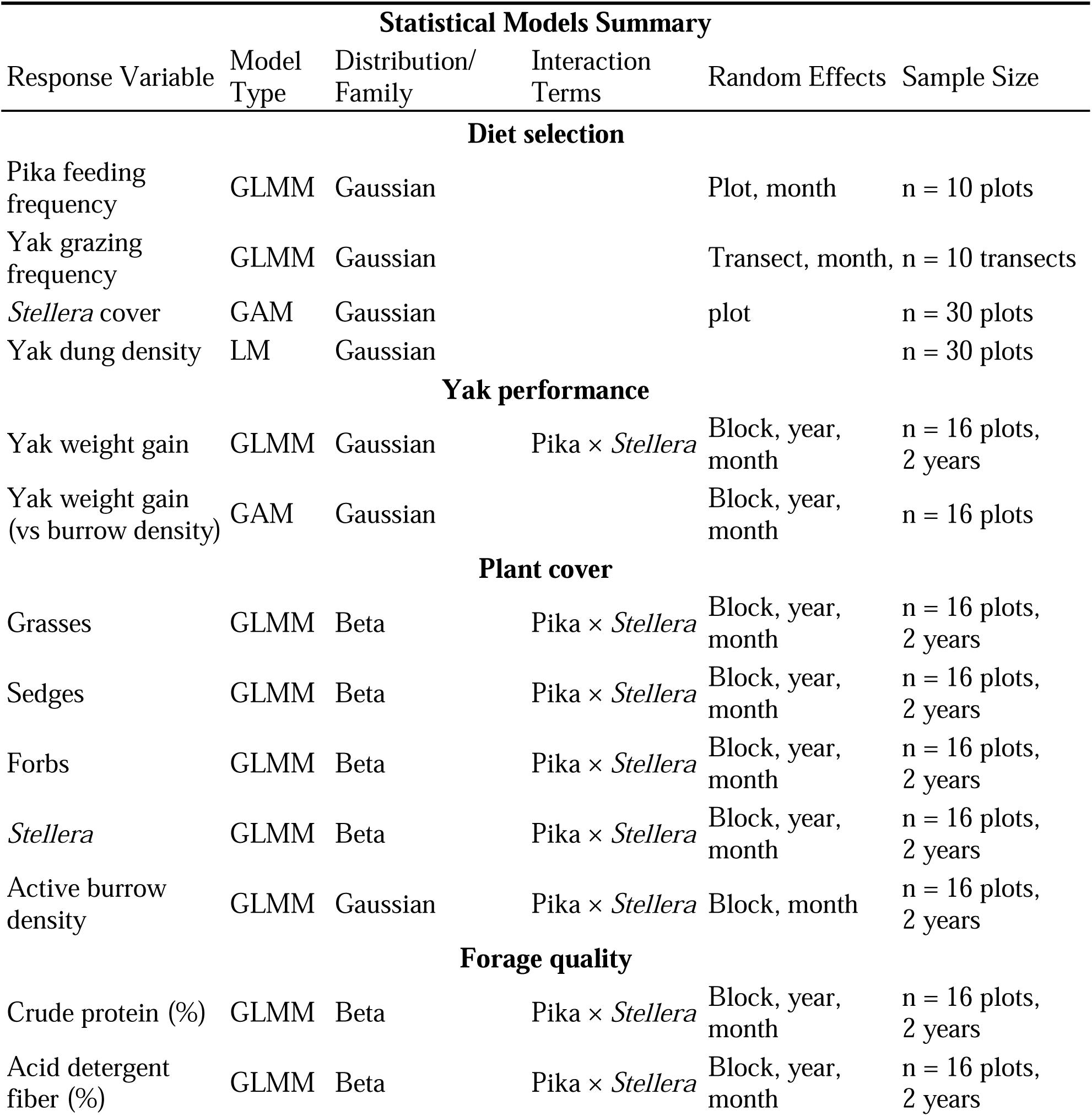
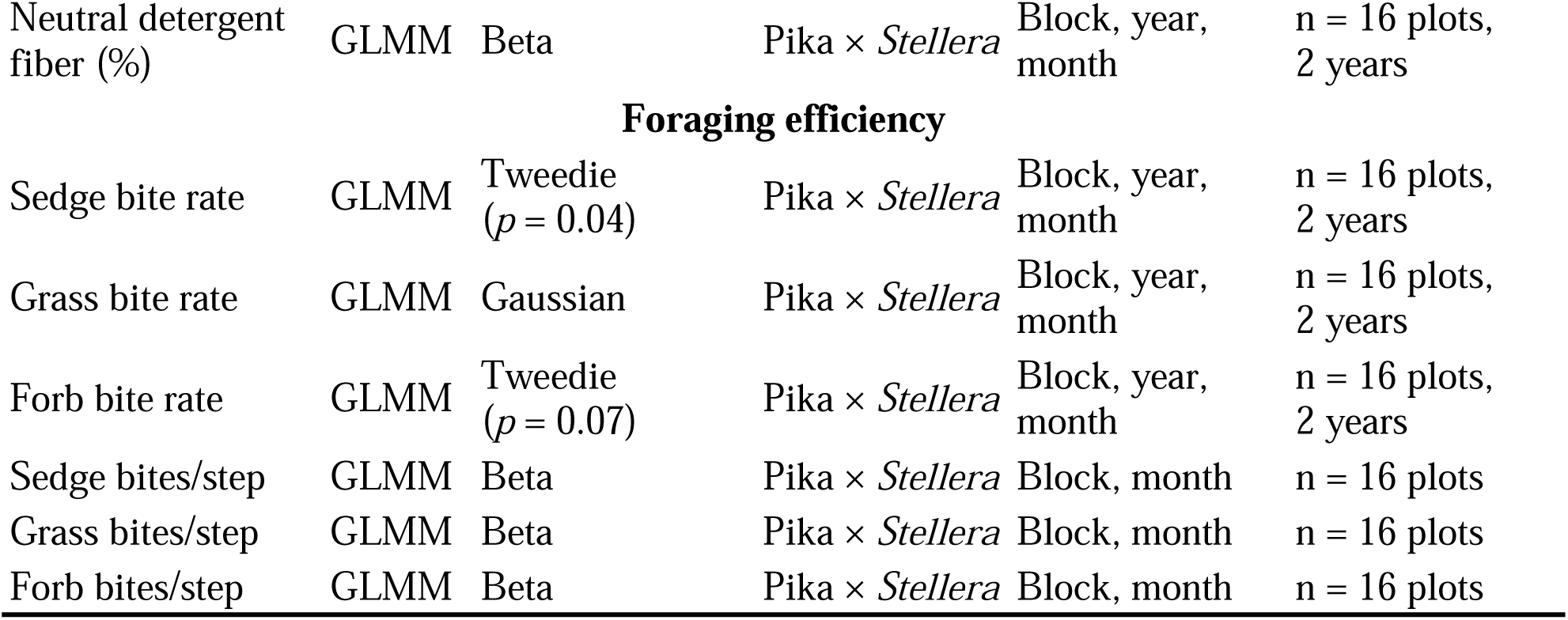
Summary of all statistical models used in the study, including response variables, model type, distribution family (with Tweedie power parameter where applicable), interaction terms, random effects structure, and sample sizes. All field experiments (2022-2023) used a 2 × 2 factorial design with pika presence/absence and *Stellera* presence/absence treatments. GLMM = Generalized Linear Mixed Model; GAM = Generalized Additive Mixed Model; LM = Linear Model. For Tweedie distributions, *p* indicates the power parameter.

